# SLX4 dampens MutSα-dependent mismatch repair

**DOI:** 10.1101/2021.10.12.464076

**Authors:** Jean-Hugues Guervilly, Marion Blin, Luisa Laureti, Emilie Baudelet, Stéphane Audebert, Pierre-Henri Gaillard

## Abstract

The tumour suppressor SLX4 plays multiple roles in the maintenance of genome stability, acting as a scaffold for structure-specific endonucleases and other DNA repair proteins. It directly interacts with the mismatch repair (MMR) protein MSH2 but the significance of this interaction remained unknown until recent findings showing that MutSβ (MSH2-MSH3) stimulates *in vitro* the SLX4-dependent Holliday junction resolvase activity. Here, we characterize the mode of interaction between SLX4 and MSH2, which relies on an M<u>SH</u>2-interacting <u>p</u>eptide (SHIP box) that drives interaction of SLX4 with both MutSβ and MutSα (MSH2-MSH6). While we show that this MSH2 binding domain is dispensable for the well-established role of SLX4 in interstrand crosslink repair, we find that it mediates inhibition of MutSα-dependent MMR by SLX4, unravelling an unanticipated function of SLX4.

## INTRODUCTION

Genetic integrity is constantly threatened by DNA lesions arising from both endogenous and exogenous origins. Cells rely on elaborate DNA repair and signalling pathways to fix or tolerate DNA damage, which determine cell survival and the level of mutagenesis. The human SLX4 protein contributes to many aspects of genome maintenance through multiple protein-protein interactions. One of its primary and best understood functions is how it acts as a nuclease scaffold that controls the XPF-ERCC1, MUS81-EME1 and SLX1 structure-specific endonucleases (SSE), directly modulating their catalytic activity and promoting their targeting to appropriate substrates (1–5). It makes key contributions to the repair of interstrand crosslinks (ICL) where it recruits XPF-ERCC1 to replication forks stalled by an ICL and stimulates the so-called unhooking of the lesion by XPF-ERCC1 (6, 7). Recruitment of SLX4 at the ICL-stalled fork is mediated by its ubiquitin binding UBZ4 domains (8–10) and while monoubiquitinated FANCD2 has been proposed to be the UBZ4 ligand that drives SLX4 recruitment, this has been under debate (7, 9–11). Nevertheless, SLX4 is part of the Fanconi pathway as underscored by the identification of rare cases of bi-allelic mutations of the *SLX4/FANCP* gene causative of Fanconi Anemia (FA) (8, 12). FA is a severe hereditary syndrome that is invariably characterized by a profound ICL hypersensitivity at the cellular level and is associated with bone marrow failure, developmental defects and cancer predisposition. In addition to promoting the endonucleolytic processing of ICLs by XPF-ERCC1, SLX4 is required for Holliday junction (HJ) resolution by MUS81-EME1 and SLX1 during the late steps of homologous recombination (HR) (13–15). SLX4 contributes to the maintenance of specific genomic loci such as telomeres and common fragile sites. Telomeric functions rely primarily on direct interaction with TRF2, which drives recruitment of SLX4 and its associated SSEs to chromosomes ends (16, 17) as well as that of SLX4IP, which was recently shown to fulfil several important functions in telomere maintenance via the alternative lengthening of telomeres pathway (18–20). SLX4 contains SUMO-interacting motifs (SIM) that also contribute to its telomeric localization and to laser-induced DNA damage (21–23) as well as to functions of SLX4 in the maintenance of common fragile sites (CFS) (21, 22) where it triggers mitotic DNA synthesis (MiDAS) (24). Recently, SLX4 was also found to prevent replication-transcription conflicts through direct interaction with the RTEL1 helicase (25).

Hence, SLX4 exerts multiple functions in genome maintenance, each mediated by one or several protein-protein interactions. Intriguingly, the mismatch repair (MMR) factor MSH2 was also identified as a binding partner of human SLX4 (3) but the functional relevance of this interaction remained unexplored until recently (26). The primary role of MMR is to correct replication errors introduced by DNA polymerases. As such, MMR deficiency is the main cause of hereditary nonpolyposis colorectal cancer (HNPCC or Lynch syndrome) (27). MSH2 acts in the early steps of MMR and is an obligate component of the heterodimeric MutSα (MSH2-MSH6) and MutSβ (MSH2-MSH3) ATPase complexes. MutSα, which is the more abundant complex, recognizes single nucleotide mismatches and small insertion/deletions (1 or 2 nt indels) while MutSβ recognizes larger indels (28). Mismatch recognition allows the recruitment and activation of the MutLα (MLH1-PMS2) endonuclease and EXO1 exonuclease that will remove the mismatch and trigger subsequent DNA repair synthesis (29–31). Noteworthy, MMR activity is responsible for the cytotoxic effect of chemotherapeutic drugs such as the alkylating agent N-methyl-N’-nitro-N-nitrosoguanidine (MNNG) or the pro-drug 6-thioguanine (6-TG), which can both induce mispairing of a methylated guanine with a newly incorporated thymidine during replication and subsequent MMR activity (32).

Immunoprecipitation of overexpressed SLX4 coupled to mass spectrometry (IP-MS) analyses suggested that it associates with MutSβ (3), MutSα (33) or both complexes (34). Noteworthy, interaction between SLX4 and MutSβ, but not MutSα, was shown to stimulate HJ resolution by the SLX4-SLX1 and SLX1-SLX4-MUS81-EME1-XPF-ERCC1 (SMX) HJ resolvase complexes *in vitro* (26). Furthermore, co-depletion of MSH3 and SLX4 did not further exacerbate the phenotypes associated with the reduced processing of recombination intermediates caused by depleting SLX4 alone, suggesting that SLX4 and MutSβ also collaborate *in vivo* (26).

In this study, we undertook a detailed analysis of the interaction between SLX4 and MSH2. We precisely characterise the SLX4-MSH2 binding interface in both proteins. We show that it involves the association between an M<u>SH</u>2-<u>i</u>nteracting <u>p</u>eptide (SHIP box) that we found in the N-terminus of SLX4 and the so-called Lever 1 domain of MSH2. Furthermore, we find that SLX4 can interact with both MutSβ and MutSα. Cellular functional analyses exploiting a CRISPR-Cas9-engineered cell line that produces an N-terminally truncated SLX4 protein reveal that whilst the SLX4-MSH2 interaction does not contribute to the function of SLX4 in ICL repair, it dampens the MutSα-dependent MMR activity *in vivo* and *in vitro*.

## MATERIAL AND METHODS

### Cell lines Generation

HeLa Flp-In T-Rex (FITo: parental cells, kindly provided by Stephen Taylor) were maintained in DMEM, 10% FBS, Pen/strep (Gibco) + 4 μg/ml Blasticidin (Invivogen). In order to knock-out *SLX4* gene in these cells, we used a CRISPR-Cas9 approach with commercially available plasmids from Santa Cruz Biotechnology (sc-404395 and sc-404395-HDR) allowing the insertion of an exogenous plasmid conferring Puromycin resistance at the endogenous *SLX4* locus. Cells were subsequently selected by 0.8 μg/ml Puromycin (InvivoGen) and individual resistant clones were isolated and tested for MMC sensitivity and Western blotting to detect SLX4. Several MMC hypersensitive clones were selected but only one (KO30) initially showed an apparent knock-out of SLX4 by Western blot while the others displayed SLX4 forms with molecular weights (MW) distinct from WT SLX4. Although KO30 also presented a lower MW form of SLX4 in subsequent experiments, this clone was further studied and complemented with Flag-HA (FHA)-tagged forms of SLX4 WT, UBZ-mutated or lacking the MSH2 binding domain (ΔMSH2bd) using the Flp-In system by co-transfecting them with pDEST-FRT-TO-FHA-SLX4 vectors and the POG44 plasmid (encoding the Flp recombinase). Recombinant clones were selected with 120 μg/ml Hygromycin (Invitrogen) and pooled to obtain a stable population maintained in medium containing Puromycin, Blasticidin and Hygromycin.

### Mutagenesis, cloning and molecular biology

Site-directed mutagenesis for the generation of SLX4^ΔMSH2bd^ was achieved using the following primers: Fwd 5’ GCAGACCCCGAGCGTTTGAGAC 3’ and Rev 5’ CAATTGTGCTGTGCGGGGTTTG 3’. pENTR1A SLX4 WT was used as a template and PCR was performed with the Advantage HD polymerase (Clontech). The PCR product was then digested by DpnI and subsequently phosphorylated and ligated using T4 PNK and T4 ligase (NEB) before transformation of *E. coli* DH5α. Clones harboured the expected loss of one AvaII restriction site and the SLX4 insert of one clone was fully sequenced. A gateway LR reaction was then performed to get the pDEST-FRT-TO-FHA and pDEST-FRT-TO-YFP expression vectors of SLX4^ΔMSH2bd^.

All the MSH2 constructs and the M453I mutant were obtained through gene synthesis or mutagenesis and cloned in pcDNA3.1(+)-N-eGFP by GenScript. These MSH2 constructs contain a N-terminal SV40 NLS to achieve nuclear localization.

Genomic DNA extraction using 4.10^6^ HeLa FITo or HeLa KO30 cells was performed using the DNeasy Blood & Tissue Kit (Qiagen). PCR was performed on 100 ng genomic DNA using PrimeStar GXL SP DNA polymerase (Takara Bio RF220Q) to check exon 3 integrity and plasmid insertion using the following primers:

Fwd: 5’ AGGAGCTGACAGAGCAGAGG 3’

Rev: 5’ TGAGGTGCTGTTGTCATGGT 3’

Fwd^PURO^: 5’ GCAACCTCCCCTTCTACGAGC 3’

### Antibodies and western blot

SDS-PAGE and western blotting were performed using a Novex NuPAGE SDS-PAGE Gel System and XCell II blot module (Invitrogen). Hybond-C Extra membrane (RPN203E) was purchased from GE healthcare. ECL Prime (RPN2236) or ECL Select (RPN2235) WB Detection Reagents were from Cytiva. A Chemidoc MP imaging system (Biorad) was used for signal detection.

Primary antibodies against SLX4 (A302-270A, A302-269A), EXO1 (A302-640A), MSH6 (A300-022A), MSH3 (A305-314A) and pS4-S8 RPA32 (A300-245A) were purchased from Bethyl laboratories. MSH2 (ab70270) and RPA32 (ab2175) antibodies were from Abcam. Anti-MSH2 (3A2 #2850), anti-MSH6 (3E1 #12988), anti-p345CHK1 (133D3 #2348) and anti-p68CHK2 (C13C1 #2197) were from Cell Signaling Technology. Anti-XPF (AM00551PU-N) was from Acris. FANCD2 (sc-20022) and PCNA (sc-56) antibodies were purchased from Santa Cruz Biotechnology. Anti-FLAG (F3165) and anti-GFP (JL-8) were from Sigma and Clontech, respectively. Anti-RPA70 (NA13) was purchased from Calbiochem.

The following secondary antibodies were purchased from Dako: goat anti-rabbit immunoglobulin G (IgG)/horseradish peroxidase (HRP) (P0448), goat anti-mouse IgG/HRP (P0447) and rabbit anti-goat IgG/HRP (P0449). To avoid the signal of the IgG used for immunoprecipitation, specific secondary antibodies were eventually used in IP/WB experiments (Veriblot, Abcam).

### Transient transfections and co-immunoprecipitation

HeLa cells were transfected with lipofectamine 2000 (Invitrogen) or PEI (gift from Mauro Modesti). If needed, overexpression of FHA-SLX4 was typically achieved with 100 ng/ml of doxycycline (Sigma). Cells were harvested 24 h after transfection and the pellet was usually frozen for at least one night at - 80°C. Frozen pellets were lysed in NETN buffer (50 mM Tris-HCl [pH=8.0], 150 mM NaCl, 1 mM EDTA, 0.5% [v/v] NP-40 supplemented with Complete EDTA-free protease inhibitor cocktail (Roche)) with rotation at 4°C before sonication and clarification by centrifugation. Immunoprecipitations (IPs) were performed overnight at 4°C with anti-FLAG M2 beads (Sigma), GFP trap (Chromotek) or anti-SLX4 (Bethyl, A302-269A and/or A302-270A) and control rabbit IgG (Cell Signaling, 2729S) coupled to dynabeads-protein G (Invitrogen). Beads were extensively washed with NETN buffer before elution in loading buffer.

For immunoprecipitation on solubilized chromatin, an adaptation of a published protocol was used (35). HeLa cells were washed in PBS and resuspended in solution A (10 mM HEPES at pH 7.9, 10 mM KCl, 1.5 mM MgCl_2_, 0.34 M sucrose, 10% glycerol, 1 mM DTT and protease inhibitors). After addition of Triton X-100 to a final concentration of 0.1%, cells were incubated on ice for 5 min before low-speed centrifugation (1300g for 4 min at 4 °C). The nuclei pellet was washed once with solution A before lysis with solution B (3 mM EDTA, 0.2 mM EGTA, 1 mM DTT and protease inhibitors) for 30 min with rotation at 4°C. After centrifugation (1700g for 4 min at 4 °C), the nuclear soluble fraction was removed and the insoluble pellet was resuspended in 1 ml Benzonase buffer (20 mM Tri-HCl [pH=8.0], 2 mM MgCl2, 20 mM NaCl with protease inhibitors) before addition of 1.2 μL of Benzonase (Sigma). After an overnight incubation at 4°C, Benzonase was added again for 4 additional hours before high speed centrifugation with the supernatant representing the solubilized chromatin further used in IP as described above.

### siRNA

Cells were transfected with the following siRNA at a concentration of 5 nM using INTERFERin (Polyplus transfection):

siLUC (CGUACGCGGAAUACUUCGA*dTdT*)

siMSH2 (AAUCUGCAGAGUGUUGUGCUU*dTdT*) [from (36)]

siSLX4-1 is a mix of two siRNA targeting the UTRs of SLX4: SLX4 UTR87 (GCACCAGGUUCAUAUGUAU*dTdT*) and SLX4 UTR7062 (GCACAAGGGCCCAGAACAA*dTdT*),

siSLX4-2 is a pool of siRNA synthetized by Dharmacon (M-014895-01-0005)

siSLX4-3 (AAACGUGAAUGAAGCAGAA*UU*) [from (3)]

siSLX4-4 (CAGATCTCAGAAATCTTCATCCAAA) is a Stealth siRNA synthetized from Invitrogen.

Unless otherwise specified, siRNAs were purchased from Eurofins MWG Operon.

### Clonogenic survival assay

Cell lines (FITo, KO30, KO30+WT, KO30+ΔMSH2bd, KO30+UBZ^mut^) were seeded at low density (450 to 500 cells) in 60 mm Petri dishes. Moderate expression of exogenous FHA-SLX4 was achieved by addition of 2 ng/ml of doxycycline throughout the entire experiment. For siRNA treatment, cells were transfected the day before at 5nM siRNA in 6 well plates before low density seeding. Genotoxic treatments with mitomycin C, melphalan, 6-thioguanine or N-methyl-N’-nitro-N-nitrosoguanidine (Sigma) were performed the next day for 24h before drug retrieval, PBS wash and addition of fresh medium. Cells were usually fixed and stained 7 to 8 days later when visible colonies could be counted with a Scan 1200 automatic colony counter (Interscience).

### Peptide pulldown

To prepare nuclear extracts for the pulldown, frozen HeLa FITo cell pellets were resuspended in solution A (10 mM HEPES at pH 7.9, 10 mM KCl, 1.5 mM MgCl_2_, 0.34 M sucrose, 10% glycerol, 1 mM DTT and protease inhibitors). After addition of Triton X-100 to a final concentration of 0.1%, cells were incubated on ice for 10 min before low-speed centrifugation (1300g for 4 min at 4 °C). The resulting nuclei pellet was washed once with solution A before lysis in extract buffer (50 mM Hepes, pH 7.3; 150 mM NaCl; 10% sucrose supplemented with protease inhibitors) and incubated with rotation for 1 h at 4 °C including one round of sonication. Pre-washed streptavidin magnetic beads (Genscript) were added for an additional 30 min and nuclear extracts were subsequently clarified by high-speed centrifugation. Protein concentration was measured using Pierce 660nm Protein Assay Reagent.

WT and mutant SLX4 peptides were synthetized by Genscript and resuspended in H_2_O. Peptides (20 μg) were immobilized on pre-washed streptavidin magnetic beads (Genscript) in TBS with 1 % BSA for 1 h at RT before washes in TBS to get rid of unbound peptides and resuspension in extract buffer without NaCl in order to get a final molarity of 110 mM NaCl for the pulldown. Beads were incubated with nuclear extracts (425 μg) for 1 h 15 at 4 °C, washed 4 times in extract buffer (110 mM NaCl) before elution in loading buffer.

### Nuclear extracts for *in vitro* MMR assay

Nuclear extracts preparation was essentially performed according to a published protocol with minor modifications (37). Briefly, 1-2 × 10^8^ HeLa FITo cells were collected, washed with PBS and resuspended in 12 ml of cold hypotonic buffer (20 mM Hepes, pH 7.3; 0.2M sucrose; 5 mM KCl; 0.5 mM MgCl_2_; 0.5 mM PMSF; 2 mM DTT and supplemented with Complete EDTA-free protease inhibitor cocktail (Roche)). Cells were pelleted at 2000 g for 5 min at 4 °C, resuspended in 4 ml of hypotonic buffer without sucrose and incubated on ice for 10 min. After 10 strokes of Dounce homogenizer (loose-fitting), the solution was centrifuged at 2000 g for 5 min at 4 °C. The nuclear pellet was lysed in 1 ml of extract buffer (50 mM Hepes, pH 7.3; 10% sucrose; 0.5 mM PMSF; 2 mM DTT and supplemented with Complete EDTA-free protease inhibitor Cocktail). NaCl was added and adjusted to 150 mM before incubation with rotation at 4 °C for 1 h. The nuclear suspension was centrifugated for 30 min at 15 000 g and supernatant was concentrated using Vivaspin 6 (10 kDa MWCO) centrifugal concentrator (Sartorius). Aliquots of concentrated nuclear extracts were frozen in liquid nitrogen and stored at -80 °C.

### Construction of the mismatched DNA substrate

For this study, we exploited a previously published plasmid construction (pLL1/2c) that contains a G/T mismatch in a *Pvu*II unique site (38). In order to introduce a nick downstream the G/T mismatch, pLL1 and pLL2c plasmids were modified by site-directed mutagenesis to introduce a *Bbv*CI restriction site 132bp far from the mismatch. The new plasmids were renamed pLL104 and pLL105. The mismatched DNA substrate was obtained following the gap-duplex method (39) using 200 μg of pLL104 and pLL105 and a 17mer oligo (GCAAGAATATTAACACG) that allows to ligate only the gap-duplex form we are interested in. Subsequently the DNA substrate was purified by CsCl/ethidium bromide gradient, recovered under UV light and resuspended in TE. Quantification on agarose gel estimated the final yield around 4-6 μg.

### *In vitro* MMR assay

To create a 5’ nick downstream the G/T mismatch, 2 μg of the DNA substrate were digested with 30U of Nb.BbvCI (from NEB) for 2 h at 37 °C. Some linear by-product was observed together with the nicked DNA due to random nicks caused by exposure to UV light. To eliminate the linear DNA we digested with 40U ExoV (from NEB) for 1 h at 37 °C. The nicked DNA substrate was finally extracted by phenol-chloroform followed by ethanol acetate precipitation and resuspended in TE to a final concentration of 40 ng/μl.

*In vitro* MMR assays were performed with minor modification as previously described (37): 80 ng of nicked DNA substrate were added to 50 μg of nuclear extract in a 50 μl final volume. MMR buffer was as followed: 0.1 mM each of four dNTPs, 20 mM Tris-HCl pH 7.6; 1.5 mM ATP; 1 mM glutathione; 5 mM MgCl_2_; 50 μg/ml BSA; the salt concentration was adjusted to 110 mM NaCl. SLX4 peptides (80.5 μM) were eventually added to the reaction. After incubation at 37 °C for 30 min, the reaction was terminated by the addition of 100 μl of stop solution (25 mM EDTA, 0.67% sodium dodecyl sulfate, and 300 μg/ml proteinase K), then incubated at 37 °C for another 15 min. DNA was extracted twice with an equal volume of phenol/chloroform and once with chloroform, precipitated by ethanol/acetate and resuspended in 10 μl H_2_O. To analyze the repair of the mismatch, the DNA substrates were digested with 10U of *ApaL*I and *Pvu*II-HF for 1 h at 37 °C (RNase was added 20min before the end of the incubation) and visualized on a 1% agarose gel. ImageJ software was used for quantification.

## RESULTS

### SLX4 associates with both MutSα and MutSβ

In agreement with previous reports (3, 33), we detected by Western blot (WB) endogenous MSH2 in SLX4 immunoprecipitates (IP) performed on whole cell extracts from HeLa cells over-expressing SLX4 (**Fig 1A**). In contrast, we were unable to detect MSH2 when we immunoprecipitated endogenous SLX4 (**Fig 1B**), suggesting that only a minor fraction of SLX4 is in complex with MSH2. Using mass spectrometry as a complementary approach, we specifically identified MSH2 and MSH6 peptides in endogenous SLX4 immunoprecipitates (**Fig S1A**), supporting endogenous SLX4-MutSα complex formation. We suspected that our difficulty to readily detect an interaction between endogenous SLX4 and MSH2 in whole cell extracts might be due to a weak and/or transient interaction and/or that it occurs preferentially on chromatin. In agreement, we successfully detected by WB MSH2, MSH6, and MSH3 in IPs of endogenous SLX4 obtained from cellular chromatin fractions (**Fig 1C**). Our results thereby demonstrate that MutSα and MutSβ complexes are *bona fide* partners of endogenous SLX4.

**Figure 1.**
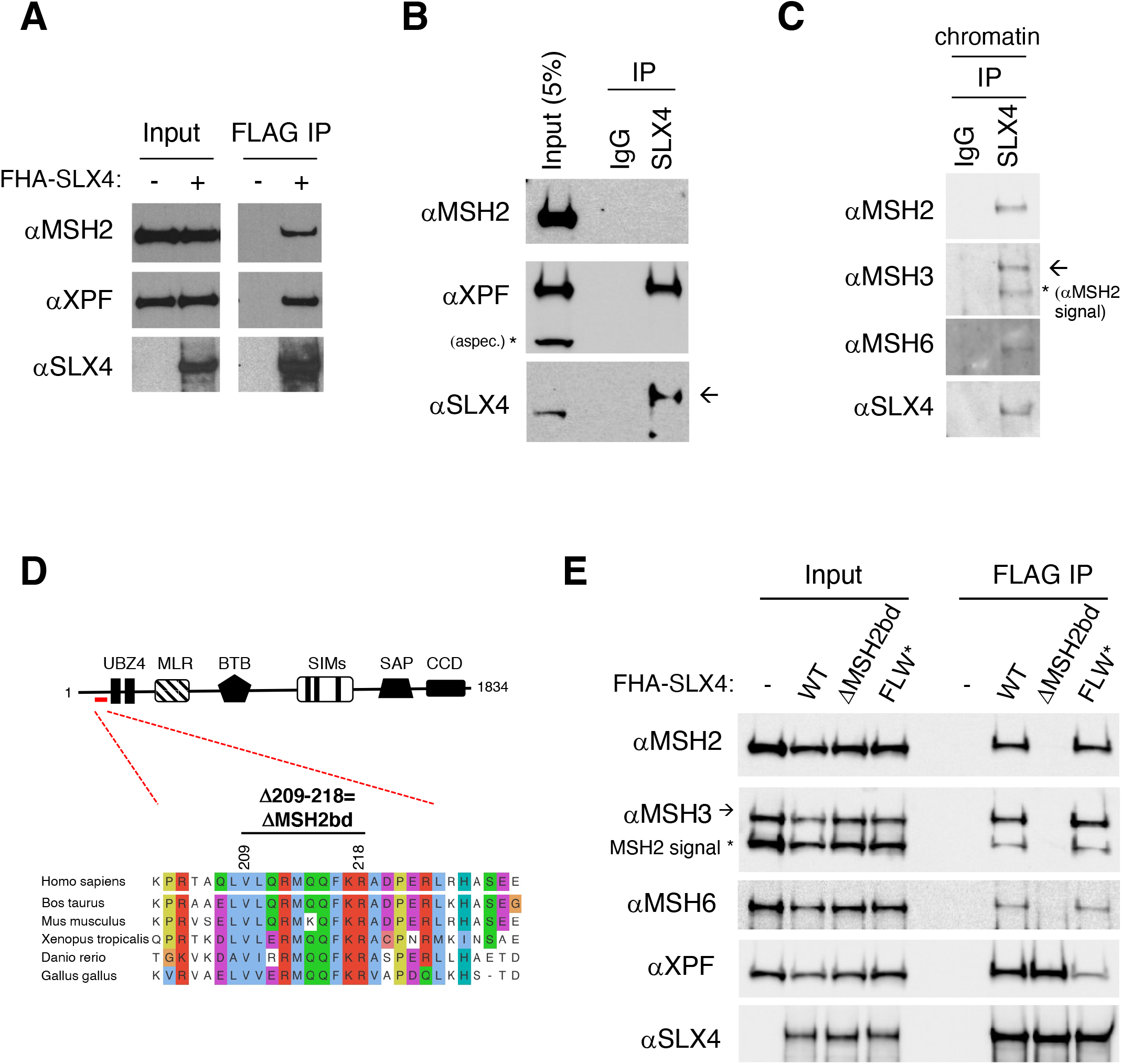
SLX4 interacts with both MutSα and MutSβ through a conserved N-terminal region. (**A**) HeLa FITo cells were transfected with FLAG-HA-SLX4 (FHA-SLX4) vector before FLAG immunoprecipitation (IP) and western blotting, which shows that MSH2 co-immunoprecipitates with overexpressed SLX4. (**B**) IP of endogenous SLX4. Co-immunoprecipitation (coIP) of MSH2 with endogenous SLX4 is barely detectable compared to experiments using overexpressed FHA-SLX4 as in A. (**C**) IP of endogenous SLX4 from a chromatin-solubilized fraction. coIP of MSH2, MSH3 and MSH6 is readily detected. (**D**) Scheme of SLX4 illustrating the location and conservation of a short domain representing the putative MSH2 binding domain (MSH2bd) deleted in SLX4^ΔMSH2bd^. Alignments were performed with ProViz (83). (**E**) HeLa FITo cells were transfected with FHA-SLX4 WT, FHA-SLX4^ΔMSH2bd^ or FHA-SLX4^FLW*^, which is deficient for XPF binding, before FLAG IP and western blotting with the indicated antibodies.

### Identification of the MSH2-binding domain in SLX4

MSH2 was initially shown to interact with a fairly large N-terminal SLX4 fragment (aa 1-669) (3). As shown in (**Fig S1B**), we tested the ability of several shorter FLAG-tagged N-terminal fragments of SLX4 to co-immunoprecipitate MSH2 when over-expressed in HeLa cells. A FLAG-SLX4^1-381aa^ fragment was sufficient to pulldown MSH2 (**Fig S1B**). In contrast, we were unable to co-immunoprecipitate MSH2 with a shorter internal FLAG-SLX4^340-580aa^ fragment, which readily interacts with XPF. This indicates that the MSH2 binding domain is located between residues 1 and 381 and that critical residues for MSH2 binding are located before residue 340. The well-described tandem UBZ4 domains of SLX4 as well as other evolutionary conserved domains of unknown function are located in between residues 1 and 380. A contribution of the UBZ motifs to MSH2 binding could be excluded as an over-expressed YFP-SLX4^UBZ^ mutant protein interacted with MSH2 as well as the YFP-SLX4 wild type control (**Fig S1C**). In contrast, a small 10 aa deletion (Δ209-218) within one of the short conserved domains found in the N-terminus of SLX4 (**Fig 1D**) totally abrogated interaction with MSH2, MSH3 and MSH6, but not XPF (**Fig 1E**). These data indicate that this conserved motif is an essential part of the MSH2 binding domain (MSH2bd) and we will refer hereafter to the SLX4 mutant lacking residues 209 to 218 as the SLX4^ΔMSH2bd^ mutant.

### Lever 1 domain of MSH2 is required for SLX4 binding

We next undertook experiments to delineate the SLX4-interacting region of MSH2. For this, we overexpressed and immunoprecipitated several deletion constructs of GFP-tagged MSH2 (**Fig S1D**). An MSH2^1-460aa^ N-terminal fragment that contains the so-called Lever 1 domain was sufficient to pull-down SLX4 but not MSH3 and MSH6. In contrast, a shorter MSH2^1-310aa^ fragment lacking the Lever 1 domain was unable to interact with SLX4 (**Fig S1D** and data not shown). Overall, our data demonstrate that the Lever 1 domain of MSH2 is critical for interaction with SLX4 and that SLX4-MSH2 complex formation can occur independently of MSH3 and MSH6. These results also strengthen the fact that it is the SLX4-MSH2 direct interaction that drives interaction with MutSα and MutSβ.

### Generation and characterization of an N-terminal truncated SLX4 cellular model

In order to determine the functional significance of the SLX4-MSH2 interaction, we took advantage of an N-terminally truncated SLX4 mutant HeLa cell line that we generated by CRISPR-Cas9-based genome editing. Using a strategy designed to knock-out *SLX4 via* the insertion of a Puromycin-resistant cassette in the first exons of the *SLX4* gene by CRISPR-Cas9 and HR, we retrieved several clones harbouring a severe MMC hypersensitivity, which is indicative of loss of SLX4 functions (**Fig S2A and S2B**). However, WB analyses with an anti-SLX4 antibody revealed bands at unexpected molecular weights (MW) in these clones, in particular a recurrent lower MW SLX4 signal (**Fig 2A, S2C and S2D**). Analysis of two of these clones (clones KO1 and KO30) showed that this signal revealed by an antibody directed against the SLX4 C-terminus is lost following SLX4 depletion by siRNA (**Fig 2A**). This strongly suggested that clones KO1 and KO30 produce an N-terminally truncated protein (termed SLX4ΔNter), consistent with our genome editing strategy that targeted the first exons of the *SLX4* gene. Using a combinatorial approach, we further characterized the origin and nature of SLX4ΔNter in KO30 cells (for details see Supplementary Results and **Fig S3A-H**). Our data indicate that KO30 cells use an alternative translation initiation site to produce a shorter SLX4^360-1834aa^ C-terminal variant that starts at Methionine 360, thus lacking the tandem UBZ4 and the MSH2bd (**Fig 2B**).

**Figure 2.**
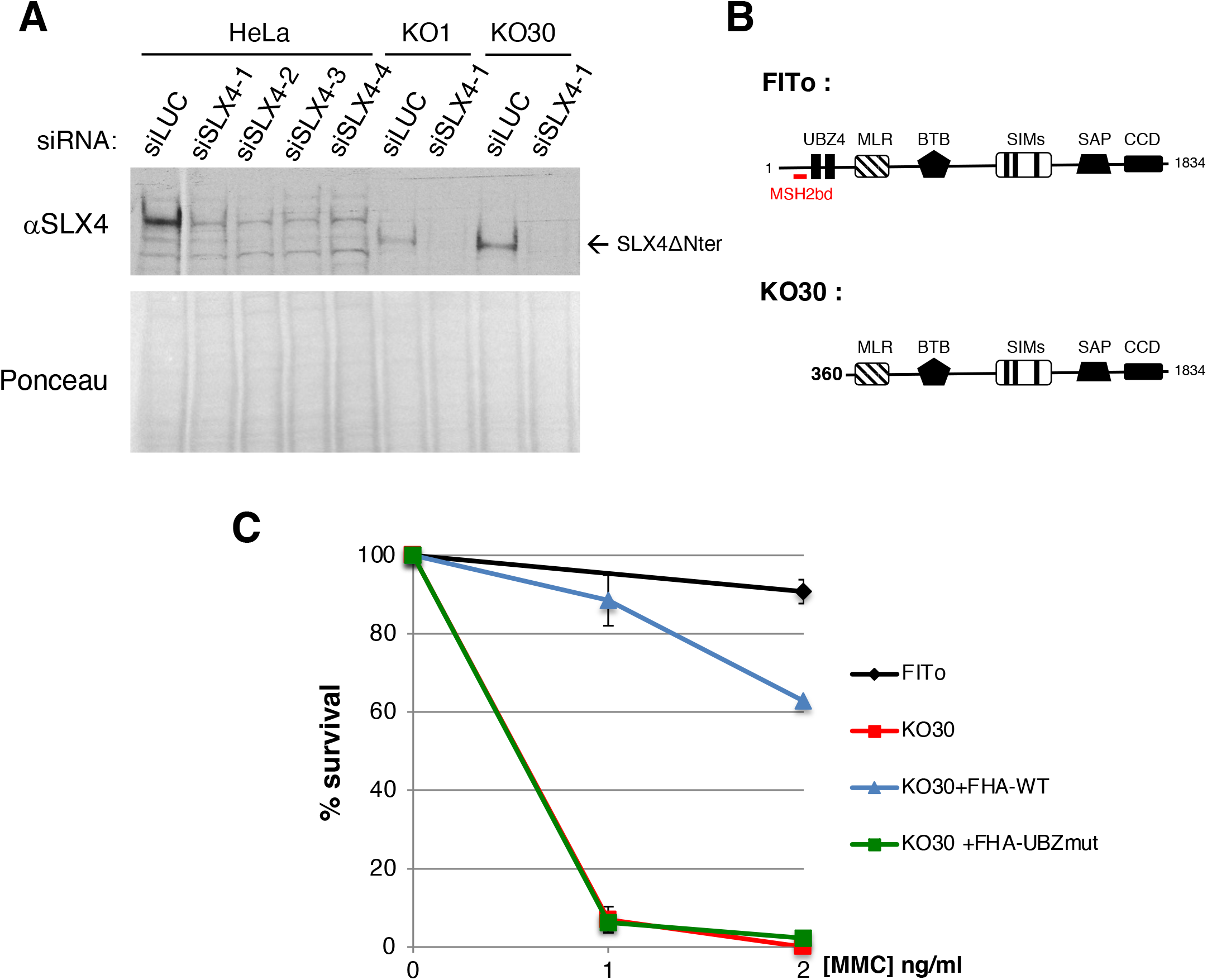
Characterization of the KO30 cell line expressing an N-terminally truncated form of SLX4. (**A**) Western blot showing that HeLa FITo KO1 and KO30 clones generated by CRISPR-Cas9 express a truncated form of SLX4 termed SLX4ΔNter (indicated by an arrow), the expression of which is sensitive to a siRNA that targets *SLX4* mRNA. (**B**) SLX4ΔNter protein starts at Methionine 360; see supplementary results for details. (**C**) Clonogenic survival assay in response to mitomycin C (MMC) of HeLa FITo, KO30 cells and KO30 cells complemented with FHA-SLX4 WT or UBZ-mutated (UBZmut). Cells were treated for 24 h with the indicated dose of MMC (n=2 to 4 experiments, mean ± SD are represented).

Loss of the SLX4 UBZ4 domains in KO30 cells is sufficient to explain their severe MMC hypersensitivity (**Fig 2C**) as the first UBZ4 motif is essential for the ICL repair function of SLX4 (10). We validated this new cellular model by complementing KO30 cells with exogenous FLAG-HA-(FHA)-tagged SLX4 WT or SLX4^UBZ^ mutant. As shown in Figure 2C, MMC hypersensitivity of KO30 cells was largely complemented by SLX4 WT but not at all by SLX4^UBZ^, establishing our KO30 cells as a worthwhile model for cellular complementation experiments aimed at studying SLX4 functions that rely on its first 359 residues.

### The SLX4-MSH2 interaction is NOT required for ICL resistance

Besides the loss of the UBZ4 domains, we could not rule out that the concomitant loss of the MSH2-binding domain in KO30 cells might also contribute to their MMC hypersensitivity since earlier studies showed that MSH2 deficiency is also associated with hypersensitivity to various ICL-inducing agents (40–42). We thus evaluated a possible interplay between MSH2 and SLX4 in ICL repair. In agreement, MSH2 depletion sensitized HeLa cells to MMC (**Fig 3A and S4A**). We thus investigated whether this function of MSH2 in ICL repair was dependent on its interaction with SLX4 by complementing KO30 cells with the SLX4^ΔMSH2bd^ mutant. Of note, this mutant appeared to be expressed in KO30 cells at lower levels than the WT protein in two independent complementation experiments (**Fig 3B and data not shown**). Nevertheless, SLX4^ΔMSH2bd^ could restore MMC resistance of KO30 cells as well as SLX4 WT (**Fig 3C**), indicating that the role of MSH2 in ICL repair is independent of SLX4. It has been suggested that MSH2 plays a more prominent role in the repair/detection of ICLs that induce a significant DNA helix distortion (43). As MMC ICLs induce a minor distortion, we assessed whether the MSH2-SLX4 interaction contributed to the repair of ICLs induced by Melphalan, which significantly distort the DNA double-helix (44). We observed similar complementation levels of the marked sensitivity of KO30 cells to Melphalan with both SLX4 WT and SLX4^ΔMSH2bd^ (**Fig 3D and S4B**). Based on these data we conclude that SLX4-MSH2 complex formation is not required for ICL repair.

**Figure 3.**
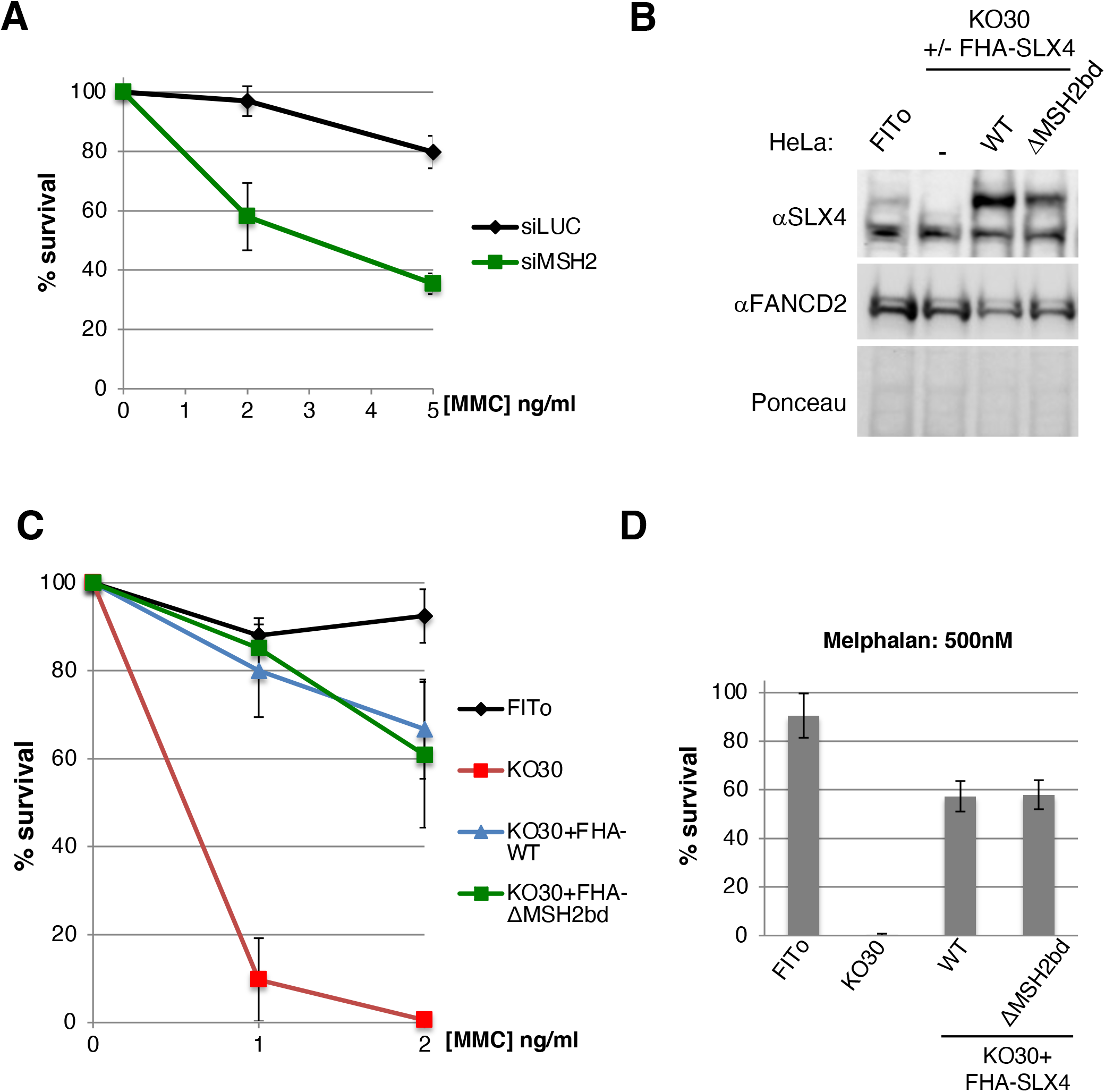
Interaction of MSH2 and SLX4 is not required for ICL repair. (**A**) Clonogenic survival assay in response to MMC of HeLa FITo cells transfected with control siRNA (siLUC) or siRNA targeting MSH2 (siMSH2) (n=3 for MMC 2 ng/ml, n=4 for MMC 5 ng/ml, mean ± SEM are represented on the graph). (**B**) Complementation of KO30 cells with FHA-SLX4 WT or SLX4^ΔMSH2bd^. Induction of exogenous SLX4 expression was achieved with 2 ng/ml of doxycycline as in (C) and (D). (**C**) Clonogenic survival assay of HeLa FITo, KO30 cells and KO30 cells complemented with FHA-SLX4 WT or SLX4^ΔMSH2bd^ in response to MMC (n=3 for MMC 1 ng/ml, n=5 or 6 for MMC 2 ng/ml, mean ± SD are represented). (**D**) Same as in (C) except that Melphalan (500 nM) was used as an alternative crosslinking agent (n=3, mean ± SD are indicated).

### The SLX4-MSH2 interaction confers resistance to 6-TG and MNNG: Is SLX4 an inhibitor of MutSα ?

As the SLX4-MSH2 interaction turned out to be dispensable for the essential role of SLX4 in ICL repair, we next tested whether SLX4 was involved in a canonical MMR function. One of the hallmarks of the majority of MMR-defective cells is their resistance to the cytotoxic effects of the long used anti-tumoral drug 6-thioguanine (6-TG). This purine analogue is a pro-drug requiring metabolic activation before incorporation into DNA during replication. Once incorporated, a methylation step generates the S^6^- methylthioguanine (S^6^-MeTG) that can readily mispair with a Thymidine (T) during the next round of replication. Perceived as a replication error, the S^6^-MeTG:T mismatch is recognized by MutSα (MSH2-MSH6) followed by the excision of the newly incorporated T. A new erroneous incorporation of a T in the daughter strand can then result in futile MMR activity leading to persistent DNA breaks and cell death (45). To assess whether SLX4-MSH2 complex formation contributes to MMR, we monitored cellular sensitivity to 6-TG of both SLX4-depleted cells and the KO30 cell line that produces N-terminally truncated SLX4 that does not bind MSH2. As expected, MSH2-depleted cells were resistant to 6-TG (**Fig 4A**). In contrast, SLX4 depletion significantly sensitized cells to 6-TG (**Fig 4A**). Remarkably, KO30 cells also displayed hypersensitivity to 6-TG, which was suppressed by SLX4 WT but absolutely not by the SLX4^ΔMSH2bd^ mutant (**Fig 4B**). If anything, complementation of KO30 with SLX4^ΔMSH2bd^ further sensitized cells to 6-TG. To substantiate our findings, we examined the response of KO30 cells complemented with SLX4 WT or SLX4^ΔMSH2bd^ after exposure to the methylating agent N-methyl-N’-nitro-N-nitrosoguanidine (MNNG). MNNG produces distinct DNA lesions but its toxicity is mainly ascribed to the generation of *O*^6^-methylguanine (^6Me^G), which can mispair with a thymidine and induce a cytotoxic MMR-dependent response in a similar way to 6-TG. As shown in Figure 4C, KO30 cells expressing SLX4^ΔMSH2bd^ were markedly more sensitive to MNNG compared to cells expressing SLX4 WT (**Fig 4C**). As MMR-mediated processing of ^6Me^G induces a checkpoint response following moderate doses of MNNG (46, 47), we examined markers of checkpoint activation in our experimental set-up. We found that phosphorylation of CHK1, CHK2 and hyperphosphorylation of RPA32 were more pronounced and persistent after MNNG treatment in KO30 cells expressing SLX4^ΔMSH2bd^ compared to those expressing SLX4 WT (**Fig 4D**). Thus, loss of interaction between SLX4 and MSH2 appears to enhance the activity of MMR. Taken together, our results demonstrate that SLX4 does not positively contribute to MMR but rather negatively impacts the repair process via its interaction with MSH2, thereby protecting cells against the MMR-mediated 6-TG and MNNG toxicity.

**Figure 4.**
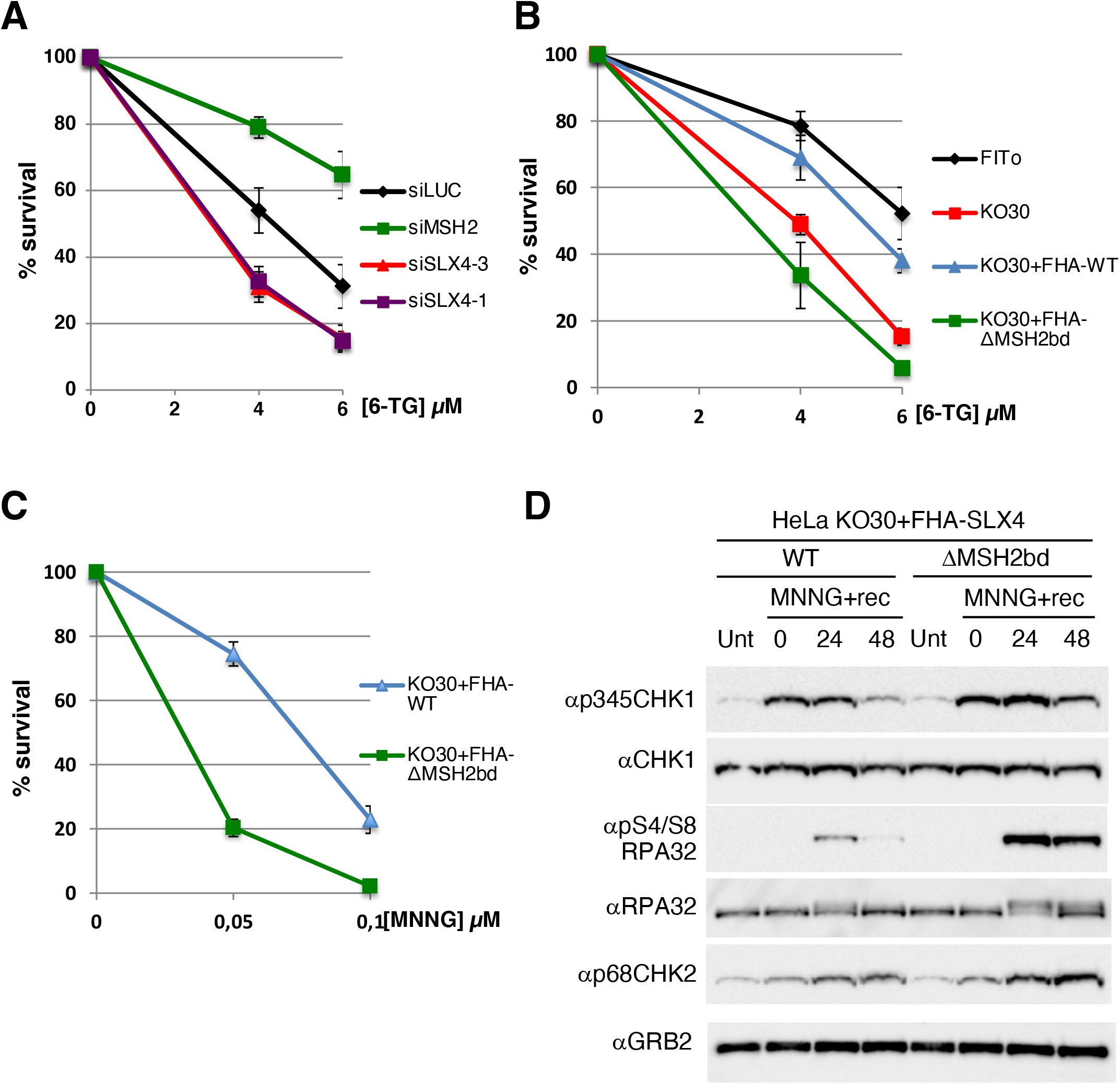
Interaction of SLX4 and MSH2 contributes to the toxicity of 6-thioguanine (6-TG) or N-methyl-N’-nitro-N-nitrosoguanidine (MNNG). (**A**) Clonogenic survival assay of HeLa FITo cells transfected with the indicated siRNA in response to a 24 h treatment with 6-TG (n=4 to 7 experiments, mean ± SEM are represented). (**B**) Clonogenic survival assay in response to 6-TG of HeLa FITo, KO30 cells and KO30 cells complemented with FHA-SLX4 WT or SLX4^ΔMSH2bd^ (n=3 to 5 experiments, mean ± SEM are represented). (**C**) Clonogenic survival assay in response to MNNG of KO30 cells complemented with FHA-SLX4 WT or SLX4^ΔMSH2bd^ (n=3 to 4 experiments, mean ± SEM are represented). (**D**) KO30 cells complemented with FHA-SLX4 WT or SLX4^ΔMSH2bd^ were treated with MNNG (0.1 μM) for 24 h before drug removal and addition of fresh medium. Cells were collected at the indicated time points of recovery (+rec) and induction of the DNA damage response was analysed by western blot.

### SLX4 negatively regulates MMR through a SHIP box-mediated interaction with MSH2

To help better understand how SLX4 may negatively impact MMR through direct binding to MSH2, we further analysed the MSH2bd *in silico*. We noticed some degree of conservation, albeit moderate, with the previously described SHIP (M<u>sh</u>2-<u>i</u>nteracting <u>p</u>eptide) boxes of *S. cerevisiae* Exo1 that drive interaction with Msh2 (48) **(Fig 5A)**. These motifs, two of which are found in the C-terminus of Exo1, have also been found in other yeast proteins (48). Mutating Methionine M470 of Msh2 into Isoleucine (Msh2^M470I^) disrupts its interaction with SHIP-box containing proteins (48). The conserved *S. cerevisiae* Msh2 M470 residue corresponds to the human MSH2 M453 residue (**Fig S5A**), which is located in an exposed helix (49) compatible with protein-protein interactions (**Fig S5B)** at the end of the Lever 1 domain of MSH2 required for binding to SLX4 **(Fig S1D**). Remarkably, as shown in **Fig 5B**, introducing the M453I mutation in MSH2 not only severely impacted interaction with EXO1, it also disrupted complex formation with SLX4 (**Fig 5B**). Our data suggest that the SHIP-box-mediated Msh2-Exo1 interaction, initially identified in *S. cerevisiae*, is conserved throughout evolution and that the MSH2bd of SLX4 is a *bona fide* SHIP box.

**Figure 5.**
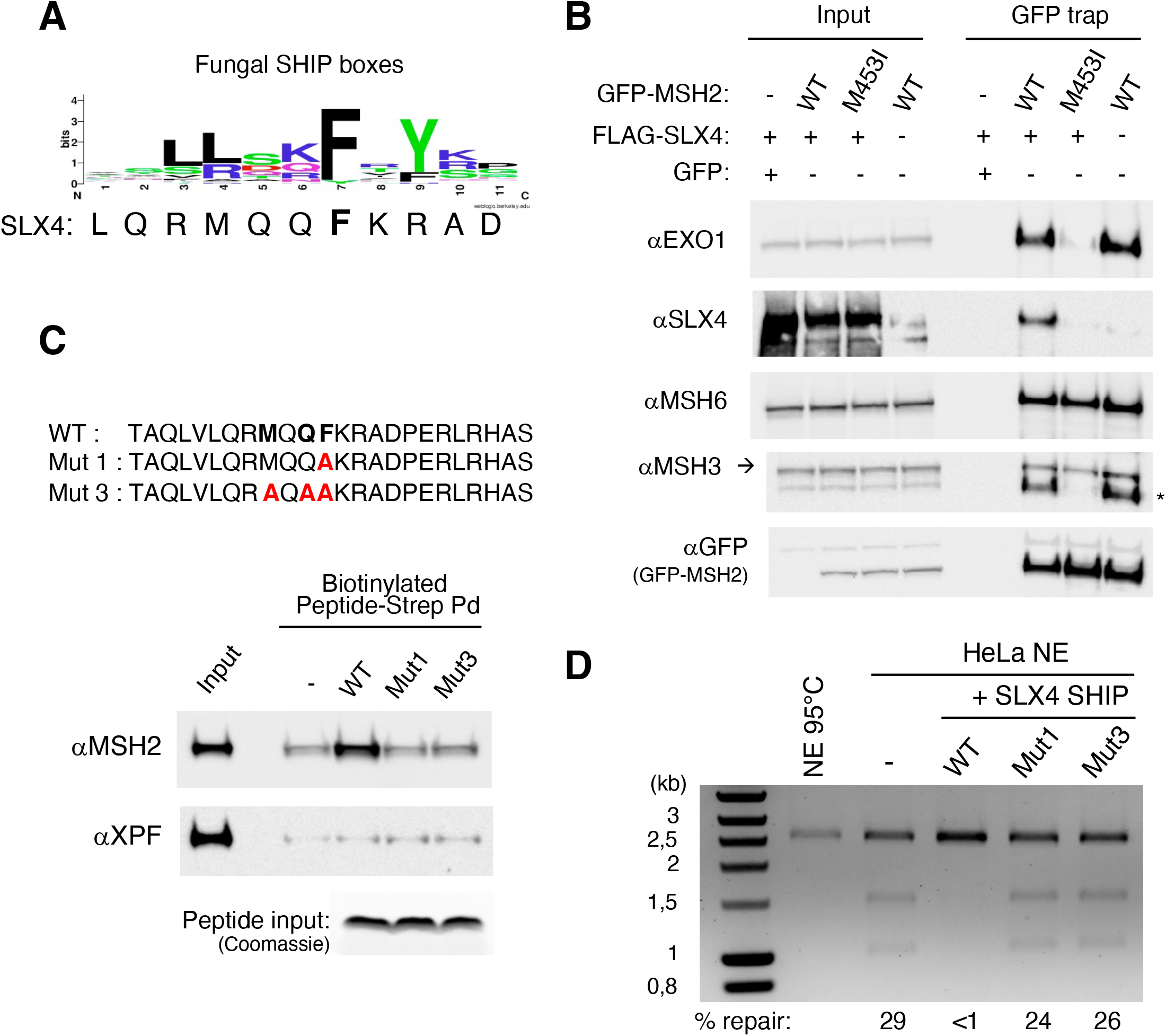
The MSH2 binding domain of SLX4 is a SHIP box that inhibits mismatch repair and antagonizes EXO1-MSH2 interaction. (**A**) Weblogo representation of multiple sequence alignments of SHIP boxes of Exo1 and Fun30 from fungal species in the *Saccharomycotina* (48, 84). (**B**) HeLa FITo cells were transfected with expression vectors coding for GFP, FLAG-SLX4, GFP-MSH2 WT and/or GFP-MSH2^M453I^, as indicated, before GFP pull-down and western blotting with the indicated antibodies. The asterisk represents the remaining signal for EXO1 when blotting for MSH3 after prior blotting for EXO1. (**C**) Peptide pull-down using biotinylated SLX4 peptides that contain a WT or mutated SHIP box, immobilized on Streptavidin-coated beads and incubated with HeLa nuclear extracts. Residues selected for mutagenesis are shown in bold in the WT sequence. (**D**) *In vitro* mismatch repair assay using a plasmid containing a G/T mismatch and a 5’ nick incubated with nuclear extracts (NE) from HeLa FITo cells as described in material and methods. Where indicated, WT or mutant SHIP peptides (80.5 μM) were added to the reaction. As a negative control, the NE was inactivated at 95°C before the reaction. DNA was purified and digested with *Apal*I and *Pvu*II. Repair of the mismatch restores the *PvuII* site and produces two bands of 1.55 kb and 1.03 kb on an agarose gel. The percentage of repair is indicated.

To definitely establish that the MSH2 binding domain is a SHIP box, we performed peptide pulldown assays using biotinylated peptides of SLX4 containing a WT or mutated SHIP box (**Fig 5C**). While endogenous MSH2 was readily pulled-down with the SLX4 WT peptide, mutation of the most conserved aromatic residue within SHIP boxes (48) (F216 in SLX4) was sufficient to disrupt MSH2 binding (**Fig 5C**) confirming that SLX4 interacts with MSH2 through a SHIP box. Finally, we investigated the ability of the SLX4 SHIP box peptide to inhibit MMR *in vitro* using a well-established assay in which a substrate containing a G-T mismatch and a 5’ nick is incubated with HeLa nuclear extracts and repair of the mismatch visualized by the restoration of a functional PvuII restriction site (**Fig 5D and S5C**) (37). Strikingly, addition of an excess of the SLX4 SHIP box peptide virtually abrogated mismatch repair while mutant peptides had barely any effect (**Fig 5D**). Overall, our data demonstrate that the SHIP box of SLX4 can inhibit MutSα-dependent MMR both *in vivo* and *in vitro* through direct interaction with MSH2.

## DISCUSSION

In this study, we have characterized the mode of interaction between SLX4 and MSH2 and investigated its functional relevance. We demonstrate that the interaction of SLX4 with MSH2 relies on a short conserved motif located upstream of the first UBZ4 domain of vertebrate SLX4 proteins. This motif resembles the small SHIP box motif previously identified in partners of *S. cerevisiae* Msh2 (48). It was previously shown that the SHIP box-mediated interaction requires the integrity of the M470 residue of yeast Msh2 (48). Importantly, we show that mutating the equivalent M453 residue at the end of the Lever 1 domain in human MSH2 (MSH2^M453I^) strongly affects the association of MSH2 with EXO1 (**Fig 5B**) thereby confirming that the SHIP box-mediated interaction with MSH2 is conserved from yeast to human, as previously anticipated (48). Importantly, MSH2^M453I^ is also strongly impaired in complex formation with SLX4 (**Fig 5B**). Furthermore, a short deletion of 10 residues within the small conserved N-terminal motif of SLX4 that is required for MSH2 binding strongly impairs its association with both MutS complexes (**Fig 1E**) and a small peptide containing that motif is sufficient to pulldown MSH2 (**Fig 5C**). Our findings thus demonstrate that SLX4 contains a *bona fide* SHIP box that drives its interaction with MSH2.

The MutSα heterodimer (MSH2-MSH6), but not MutSβ (MSH2-MSH3), mediates the cytotoxicity of 6-TG and MNNG (50–53). The underlying mechanism is likely to involve futile cycles of MMR of meG-T mispairs (54), resulting in cell death. A “direct signalling” model has also been proposed where MMR proteins directly activate ATR, without the need of processing the mismatch (55–57). These two models probably stand right and are not necessarily exclusive. In contrast to MSH2-depleted cells that gained expected 6-TG resistance, SLX4-depleted cells turned out to be hypersensitive to 6-TG (**Fig 4**). In agreement with our findings, an unbiased genome-wide CRISPR-Cas9 screen recently identified SLX4 as a determinant of MNNG resistance (58). Moreover, we demonstrated that the pro-survival role of SLX4 in response to 6-TG or MNNG is totally dependent on its interaction with MSH2 (**Fig 4**). This strongly suggests that SLX4 reduces MutSα-dependent MMR cytotoxic activity in response to these drugs. Besides inhibition of MMR, an alternative explanation for how SLX4 mediates 6-TG and MNNG resistance could have been through the resolution of HR intermediates formed after MutSα-dependent MMR. In line with this, cells lacking BRCA2 or RAD51 paralogs were previously shown to be hypersensitive to 6-TG or MNNG (59–62) and MutSΔ stimulates *in vitro* the processing of HR intermediates by the SMX complex (26). However, MSH3-deficient cells are not sensitive to 6-TG nor MNNG making this an unlikely explanation (50–53).

How exactly SLX4 inhibits MutSα-dependent MMR is currently unclear. The identification of a SHIP box in SLX4 that mediates its interaction with MSH2 raises the possibility of a competition with other SHIP box containing proteins. Indeed, yeast Exo1 and Fun30 positively contribute to MMR via their SHIP boxes (48). In human cells, other proteins strongly suspected to interact with MSH2 via a SHIP box such as WDHD1/AND1/hCTF4, SMARCAD1 (Fun30 homolog) or MCM9 (48) are also positive regulators of MMR (63–66). Since SLX4 represents the first example of a SHIP box-containing protein that exerts an inhibition of MMR, as judged by 6-TG and MNNG toxicity analyses, it is tempting to speculate that it does so by competing with SHIP box-containing positive contributors. Amongst these, EXO1 is a strong candidate as it contributes to the cytotoxicity of 6-TG (67) and MNNG (68, 69). In line with this, transient overexpression of FLAG-SLX4 decreased EXO1 interaction with GFP-MSH2 **(Fig 5B**). Moreover, we found that the interaction of GFP-MSH2 with endogenous EXO1 was slightly increased in KO30 cells compared to parental HeLa FITo cells (**Fig S5D**). Similarly, enhanced interaction of EXO1 with GFP-MSH2 was observed in KO30 cells complemented with SLX4^ΔMSH2bd^ compared to cells complemented with SLX4 WT (**Fig S5E**). Based on these observations we propose that SLX4 reduces the toxicity of 6-TG and MNNG, at least in part, by competing with EXO1 for MSH2 binding thereby limiting the rate of MMR and the associated futile cycle. In absence of SLX4 or when SLX4 cannot bind MSH2, an increased number of MMR transactions are taken to completion resulting in overall enhanced MMR activity and subsequent toxicity in response to 6-TG and MNNG. Bringing further support to this model, we showed that adding an excess of SLX4 SHIP box peptides to HeLa cell nuclear extracts abrogated 5’-directed repair of a G-T mismatch *in vitro* (**Fig 5D**). This peptide-based approach was initially used to demonstrate an early role of PCNA in MMR (preceding DNA resynthesis) using a p21 peptide containing a PCNA interacting protein (PIP)-box (70). The *S. cerevisiae* Exo1 SHIP1 box was also shown to inhibit MMR *in vitro* (48). Although these *in vitro* results obtained with the SLX4 SHIP box peptide are striking, the mechanism by which SLX4 mitigates MMR *in vivo* is certainly more complex. In line with this, we could not detect a significant difference in MMR activity between nuclear extracts from KO30+SLX4 WT versus KO30+SLX4^ΔMSH2bd^ cells (data not shown). A likely explanation is that our peptide-based experiment uses a vast excess of the SLX4 SHIP box peptide over EXO1 that will disrupt the EXO1-MSH2 interaction whereas the concentration of SLX4 in nuclear extracts, even in those derived from cells over-producing SLX4, might not be high enough to sufficiently compete out EXO1. *In vivo*, dampening of MMR by SLX4 must be tightly regulated and may rely on actively driven high concentrations of SLX4 at specific genomic locations or in the vicinity of replication forks that will locally impact MutSα-dependent repair. Furthermore, our data suggest that SLX4-MSH2 complex formation may preferentially occur on chromatin (**Fig 1C**) and there are precedents for chromatin-dependent regulation of MMR (64, 65, 71, 72), such as the targeting of MutSα to the epigenetic mark H3K36me3 (71, 72). Hence, recapitulating SLX4-driven MMR inhibition in cell-free extracts might prove particularly challenging. Alternatively, SLX4 may dampen MMR preferentially in the context of mismatches that contain a modified base such as meG-T induced by 6-TG or MNNG. However, SLX4 has been detected in an unbiased proteomic study that searched for proteins that preferentially associate with a plasmid containing an A/C mismatch in Xenopus nucleoplasmic extracts (64), suggesting that recruitment of SLX4 by MutSα and modulation of MMR is not limited to mismatches containing a modified base.

Given the primordial role of MMR repair in maintaining genome stability, it may appear surprising at first glance that evolution has selected SLX4-driven MMR inhibition. However, this is not unprecedented and other mechanisms of MMR inhibition have been described. As mentioned above, p21 inhibits PCNA function in MMR (70). Furthermore, the deacetylase HDAC6 also negatively regulates MMR by promoting MSH2 degradation (73) and preventing MLH1 binding to MutSα (74) while CAF1-mediated replication coupled chromatin assembly was proposed to limit the extent of MMR driven degradation of the nascent strand (75, 76). Sequestration of MLH1 by FAN1 has recently been reported to reduce MutSβ -dependent MMR activity and prevent the expansion of CAG repeats (77). Inhibition of MMR has also been reported in pathological situations. For example, nuclear EGFR that is associated with poor outcome in various cancers (78), phosphorylates PCNA on tyrosine 211 and inhibits MMR, presumably by weakening the interaction of phosphorylated PCNA with MutSα and MutSΔ (79). Overexpression of HORMAD1, a meiosis-specific protein aberrantly expressed in various cancers, was also shown to inhibit MMR through the cytosolic sequestration of the MCM9 helicase (80) that normally stimulates the chromatin recruitment of MLH1 and contributes to MMR *in vivo* (66, 80). Future research will help determine in which circumstances inhibiting this crucial genome maintenance pathway is beneficial. MMR dampening by SLX4 might prove particularly relevant during meiotic recombination between homologs. In vegetative cells, MMR dampening by SLX4 may represent an important tolerance mechanism that tempers the MMR-mediated toxicity of O6-methylguanines DNA lesions that have escaped repair by the O6-methylguanine DNA methyltransferase (MGMT). As such, tumours associated with loss or low levels of SLX4 and/or SLX4 mutations that abrogate SLX4-MSH2 complex formation may be good responders to MGMT inhibitors.

Besides characterizing a new functional interaction between SLX4 and MutSα-dependent MMR, we have generated and established the KO30 cell line as a novel mutant cellular model that expresses an N-terminally truncated variant of SLX4 (SLX4ΔNter) that starts at Met360. The recovery of such cells from experiments initially aimed at knocking-out *SLX4*, suggests that SLX4 functions critical for the viability of HeLa cells are harboured by parts of SLX4 that are downstream of the N-terminal truncation. If so, generating a *bona fide* SLX4 fully knocked-out HeLa cell line might prove challenging, if not impossible, without the recovery of hypomorphic SLX4 mutant clones that display profound ICL hypersensitivity while producing shorter C-terminal SLX4 variants (**Fig 2 and Fig S2**). In line with this, it was previously reported that knocking-out SLX4 in the chicken tumoral DT40 cell line is lethal and that this does not involve the loss of SLX4 ICL repair functions (9), raising the possibility that SLX4 may also be essential in human cancer cell lines (1). The KO30 cell line represents a valuable model to better characterize the functions of the SLX4^1-359aa^ N-terminal region, including those involved in ICL repair. We believe that the acute ICL hypersensitivity of KO30 cells is primarily due to the loss of the UBZ4 motifs and not to impaired SLX4-MSH2 complex formation, as it is corrected by SLX4^ΔMSH2bd^ just as well as by WT SLX4 (**Fig 3**). The lack of contribution of SLX4-MSH2 to cellular resistance to crosslinking agents may come as a surprise considering that both SLX4 and MSH2 contribute to ICL repair and the known interplay between FA proteins and MMR proteins in ICL repair (41, 81). However, MSH2 has been found to promote monoubiquitination of FANCD2 (40, 41) whereas current evidence points rather to a role of SLX4 downstream of FANCD2 monoubiquitination (6, 7, 9). Interestingly, while some MMR factors contribute to ICL repair, several FA proteins have been suggested to positively contribute to MMR (41, 82). Therefore, our findings that SLX4 (FANCP) instead negatively controls MMR shed new light on the functional ties between FA and MMR and suggest that investigating the multiple connections between FA and MMR proteins is a worthwhile line of research that can unravel unexpected findings.

In conclusion, our study unravels an unexpected function of SLX4 that involves MMR dampening driven by SLX4-MutSα complex formation. It provides a detailed mapping and functional analysis of the SLX4-MSH2 interaction yielding important structural insight with the identification of a conserved SHIP box within the SLX4 N-terminus. By showing that SLX4-MSH2 complex formation relies on the association of the SLX4 SHIP box with the Lever 1 domain of MSH2 and that it follows the same principles than EXO1-MSH2 complex formation, our findings suggest that SLX4 negatively interferes with MMR by competing with other SHIP box containing MMR activators. Future research will be needed to better understand in which context inhibition of MMR by SLX4 is important for the maintenance of genome stability and to assess the functional ties that we suspect must exist between this mechanism and cancer biology.

## ACKNOWLEDGEMENT

We are grateful to Mauro Modesti, Christophe Lachaud and Vincent Pages for sharing reagents. We thank our colleagues of the Gaillard team and the CRCM 3R community for stimulating discussions and express our gratitude to Emmanuelle Despras and Bertrand Llorente for their critical reading of the manuscript.

## Supplementary information for

### SUPPLEMENTARY RESULTS

#### Molecular characterization of SLX4 ΔNter in KO30 cells

We selected the HeLa KO clone number 30 (KO30) for further studies based on the initial apparent absence of SLX4 expression as judged by WB (**Fig S2D**). However, a band of lower molecular weight (MW) became detectable by a C-terminal specific SLX4 antibody at later passages of HeLa KO30, as in other clones (**Fig 2A, Fig S2C and S2D**). This suggested that cells producing an N-terminally truncated SLX4 protein (SLX4ΔNter), which might confer a growth advantage, are selected through successive passages (**Fig 2A**). Importantly, the SLX4ΔNter band was lost following siRNA-mediated depletion of SLX4 (**Fig 2A**) while it was specifically detected in SLX4 immunoprecipitates (IPs) from HeLa KO30 cell extracts (**Fig S3A**). Of note, an aspecific band that runs at the same position than SLX4Δ Nter was sometimes detected in whole cell extracts of parental HeLa FITo cells but not in SLX4 IPs (**Fig S3A**). The N-terminally truncated nature of the SLX4 specie detected in HeLa KO30 cells was further confirmed by IP/MS analyses of SLX4 IPs. In contrast to SLX4 IPs from the parental FITo, SLX4 IPs from HeLa KO30 cells were systematically devoid of peptides spanning the N-terminus of SLX4 (**Fig S3B**).

PCR analysis of genomic DNA from HeLa KO30 cells revealed the disruption of *SLX4* exon 3 by the insertion of the HR-plasmid (**Fig S3C and S3D**). The production of SLX4ΔNter in these cells suggested that an alternative *SLX4* mRNA that contains an alternative Translation Initiation Site (TIS) is transcribed from a start site located downstream of the plasmid insertion site. In line with this, we noticed that several alternative Transcription Start Sites (TSS) have been identified within intron 3 and at the beginning of exon 4 using TSS-seq (DataBase of Transcriptional Start Sites https://dbtss.hgc.jp/) in various cell lines (**Fig S3E**). Furthermore, our *in silico* analyses of sequences located downstream of these alternative TSS, identified Methionine 360 (Met360) as a good candidate for an alternative TIS (altTIS) that matches the Kozak sequence consensus (**Fig S3F**) (1). Furthermore, Met360 is the first aa of a tryptic peptide (**M**EVGPQLLLQAVR) that we reproducibly detected (n=14) in SLX4 IPs from HeLa FITo cells but never in SLX4 IPs from HeLa KO30 cells (**Fig S3G**). We surmised that this might be due to a post-translational modification (PTM) of Met360 in HeLa KO30 cells that blurs MS/MS analysis and peptide identification. In line with this, an estimated 97,5% of initiator Methionines (iMet) that are immediately followed by a Glutamate residue, as is Met360, are N-terminally acetylated (2). To establish whether Met360 might be acetylated in HeLa KO30 cells but not in HeLa FITo parental cells, we took into account for peptide mass calculation the possible acetylation of Methionine residues in IP/MS analyses of SLX4 IPs from nuclear extracts derived from both cell lines. As shown in **Figure S3H**, this enabled us to detect a Met-acetylated peptide (Ac-**M**EVGPQLLLQAVR) specifically in SLX4 IPs from HeLa KO30 cells, which demonstrates that Met360 serves as the iMet of SLX4ΔNter. Our data therefore unambiguously show that SLX4ΔNter is a shorter version of SLX4 (aa360-1834) produced in HeLa KO30 cells that lacks the MSH2-binding SHIP box, identified in this study, and the well characterized tandem UBZ4 ubiquitin binding zinc fingers (**Fig 2B**).

### SUPPLEMENTARY METHODS

#### Mass spectrometry analysis

##### Interactome analysis

Immunoprecipitated proteins that co-purified with SLX4 were eluted from the beads with LDS (Lithium dodecyl sulfate) sample buffer were loaded on NuPAGE 4-12% Bis-Tris acrylamide gels (Life Technologies) to stack proteins in a single band. Following staining with Imperial Blue (Thermo Fisher Scientific), protein bands were excised from the gel and gel pieces were submitted to in-gel trypsin digestion following cysteines reduction and alkylation (3). Peptides were extracted from the gel and vaccuum dried. Samples were reconstituted with 0.1% trifluoroacetic acid in 4% acetonitrile and analysed by liquid chromatography (LC)-tandem mass spectrometry (MS/MS) using an Orbitrap Fusion Lumos Tribrid Mass Spectrometer (Thermo Fisher Scientific, San Jose, CA) online with a nanoRSLC Ultimate 3000 chromatography system Thermo Fisher Scientific, Sunnyvale, CA). Peptides were separated on a Thermo Scientific Acclaim PepMap RSLC C18 column (2μm, 100A, 75 μm x 50 cm). For peptide ionization in the EASY-Spray nanosource in front of the Orbitrap Fusion Lumos Tribrid Mass Spectrometer, spray voltage was set at 2.2 kV and the capillary temperature at 275 °C. The Orbitrap Lumos was used in data dependent mode to switch consistently between MS and MS/MS. Time between Masters Scans was set to 3 seconds. MS spectra were acquired with the Orbitrap in the range of m/z 400-1600 at a FWHM resolution of 120 000 measured at 400 m/z. AGC target was set at 4.0e5 with a 50 ms Maximum Injection Time. For internal mass calibration the 445.120025 ions was used as lock mass. The more abundant precursor ions were selected and collision-induced dissociation fragmentation was performed in the ion trap to have maximum sensitivity and yield a maximum amount of MS/MS data. Number of precursor ions was automatically defined along run in 3s windows using the “Inject Ions for All Available parallelizable time option” with a maximum injection time of 300 ms. The signal threshold for an MS/MS event was set to 5000 counts. Charge state screening was enabled to exclude precursors with 0 and 1 charge states. Dynamic exclusion was enabled with a repeat count of 1 and duration of 60 s.

Raw files generated from mass spectrometry analysis were processed with Proteome Discoverer 1.4.1.14 (Thermo fisher Scientific) to search against the human protein proteome of the swissprot database (20,368 entries, extracted from Uniprot on november 2019). Database search with Mascot were done using the following settings: a maximum of two trypsin miss cleavage allowed, methionine oxidation and protein N-terminus acetylation as variable modifications and cysteine carbamidomethylation as fixed modification. A peptide mass tolerance of 6 ppm and a fragment mass tolerance of 0.8 Da were allowed for search analysis. Only peptides with high stringency Mascot scores were selected for protein identification. False discovery rate was set to 1% for protein identification.

##### Identification of the N-terminus of SLX4ΔNter

To identify the amino-terminal end of SLX4 protein, SLX4 was first immunoprecipitated from FITo (WT SLX4) or KO30 (SLX4ΔNter) nuclear extracts. Proteins were separated on NuPAGE 4-12% Bis-Tris acrylamide gels (Life Technologies) and following imperial blue staining, the upper part of the gel corresponding to proteins between MW 150 and 300 kDa was cut in 4 separate bands (respectively bands 1 to 4 for WT and 5 to 8 for KO30 extracts). Each band was digested as previously described and analyzed by liquid chromatography (LC)-tandem MS (MS/MS) using a Q Exactive Plus Hybrid Quadrupole-Orbitrap online with a nanoLC Ultimate 3000 chromatography system (Thermo Fisher Scientific™, San Jose, CA). 5 microliters corresponding to 33 % of digested protein were injected on the system. After pre-concentration and washing of the sample on a Acclaim PepMap 100 column (C18, 2 cm × 100 μm i.d. 100 A pore size, 5 μm particle size), peptides were separated on a LC EASY-Spray column (C18, 50 cm × 75 μm i.d., 100 A, 2 μm, 100A particle size) at a flow rate of 300 nL/min with a two steps linear gradient (2-20% acetonitrile/H20; 0.1 % formic acid for 40 min and 20-40% acetonitrile/H20; 0.1 % formic acid for 10 min). For peptides ionization in the EASYSpray source, spray voltage was set at 1.9 kV and the capillary temperature at 250 °C. All samples were measured in a data dependent acquisition mode. Each run was preceded by a blank MS run in order to monitor system background. The peptide masses were measured in a survey full scan (scan range 375-1500 m/z, with 70 K FWHM resolution at m/z=400, target AGC value of 3.00×10^6^ and maximum injection time of 100 ms). Following the high-resolution full scan in the Orbitrap, the 10 most intense data-dependent precursor ions were successively fragmented in HCD cell and measured in Orbitrap (normalized collision energy of 27 %, activation time of 10 ms, target AGC value of 1.00×10^5^, intensity threshold 1.00×10^4^ maximum injection time 100 ms, isolation window 2 m/z, 17.5 K FWHM resolution, scan range 200 to 2000 m/z). Dynamic exclusion was implemented with a repeat count of 1 and exclusion duration of 10 s.

Raw files generated from mass spectrometry analysis were processed with Proteome Discoverer 1.4.1.14 (Thermo fisher Scientific) to search against the human protein proteome of the swissprot database (20,368 entries, extracted from Uniprot on november 2019) modified by the addition of 85 SLX4 sequences. The Q8IY92 uniprot entry corresponding to the entire sequence 1-1834 of the SLX4 protein was used to create and add artificial 85 different SLX4 sequences corresponding to amino-terminal truncated proteins deleted from 301 to 386 first amino-acids, each sequence differing by the incremental deletion of 1 amino-acid. First sequence named Q8IY92-302 corresponds for example to 302N-N1834 SLX4 sequence and Q8IY92-387 to 387F-N1834 SLX4 sequence. Database search with Mascot were done using the following settings: a maximum of two trypsin miss cleavage allowed, methionine oxidation and protein N-terminus acetylation as variable modifications and cysteine carbamidomethylation as fixed modification. A peptide mass tolerance of 10 ppm and a fragment mass tolerance of 0.8 Da were allowed for search analysis. Only peptides with high stringency Mascot scores were selected for protein identification. False discovery rate was set to 1% for protein identification. To compare SLX4 sequence coverage and identify N-terminal sequence for both WT and truncated form of SLX4, one search of raws corresponding to bands 1 to 4 (WT) was compared to corresponding search of raws 5 to 8 from N-terminally truncated SLX4 (KO30).

### SUPPLEMENTARY FIGURE LEGENDS

**Figure S1.**
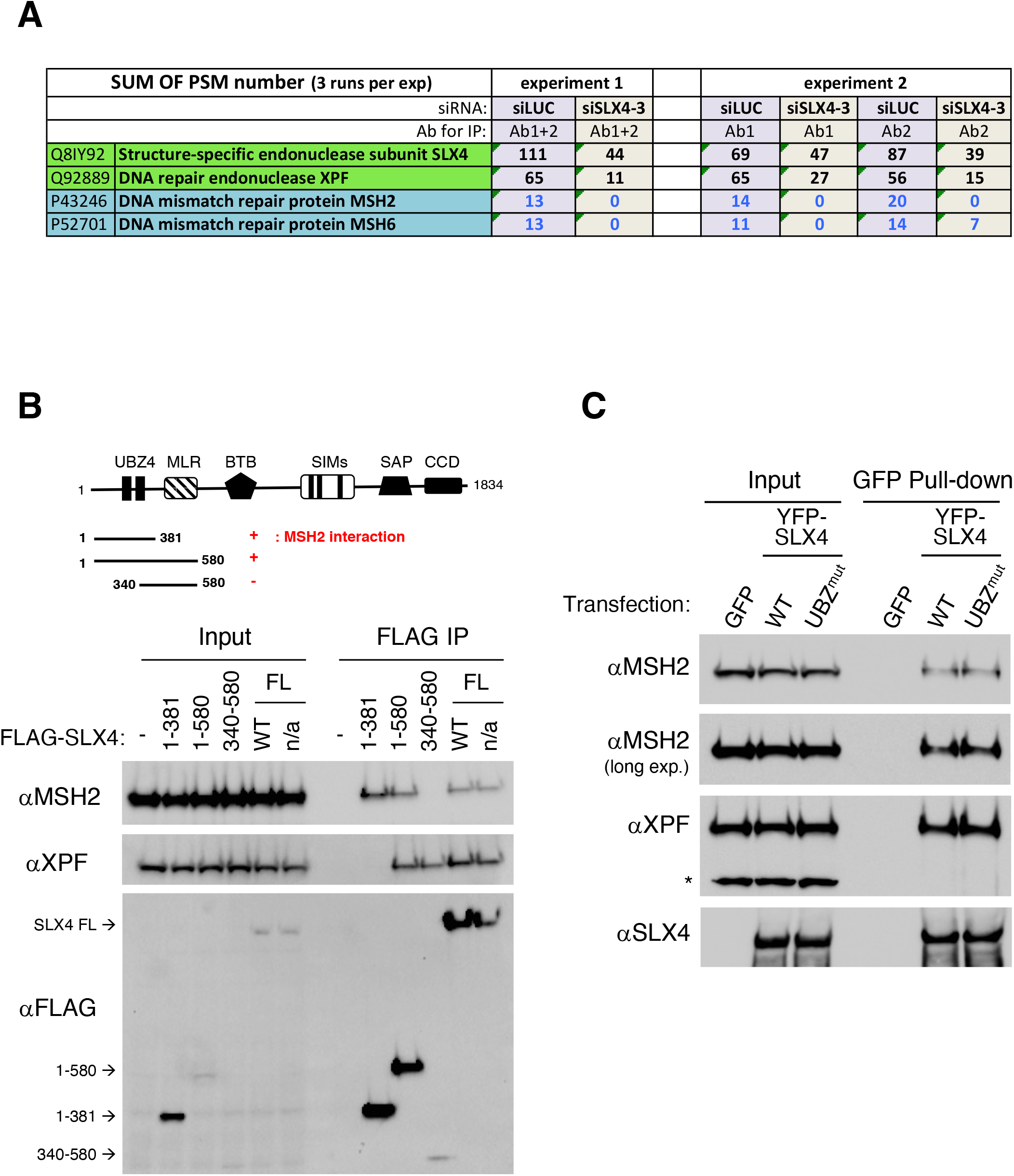
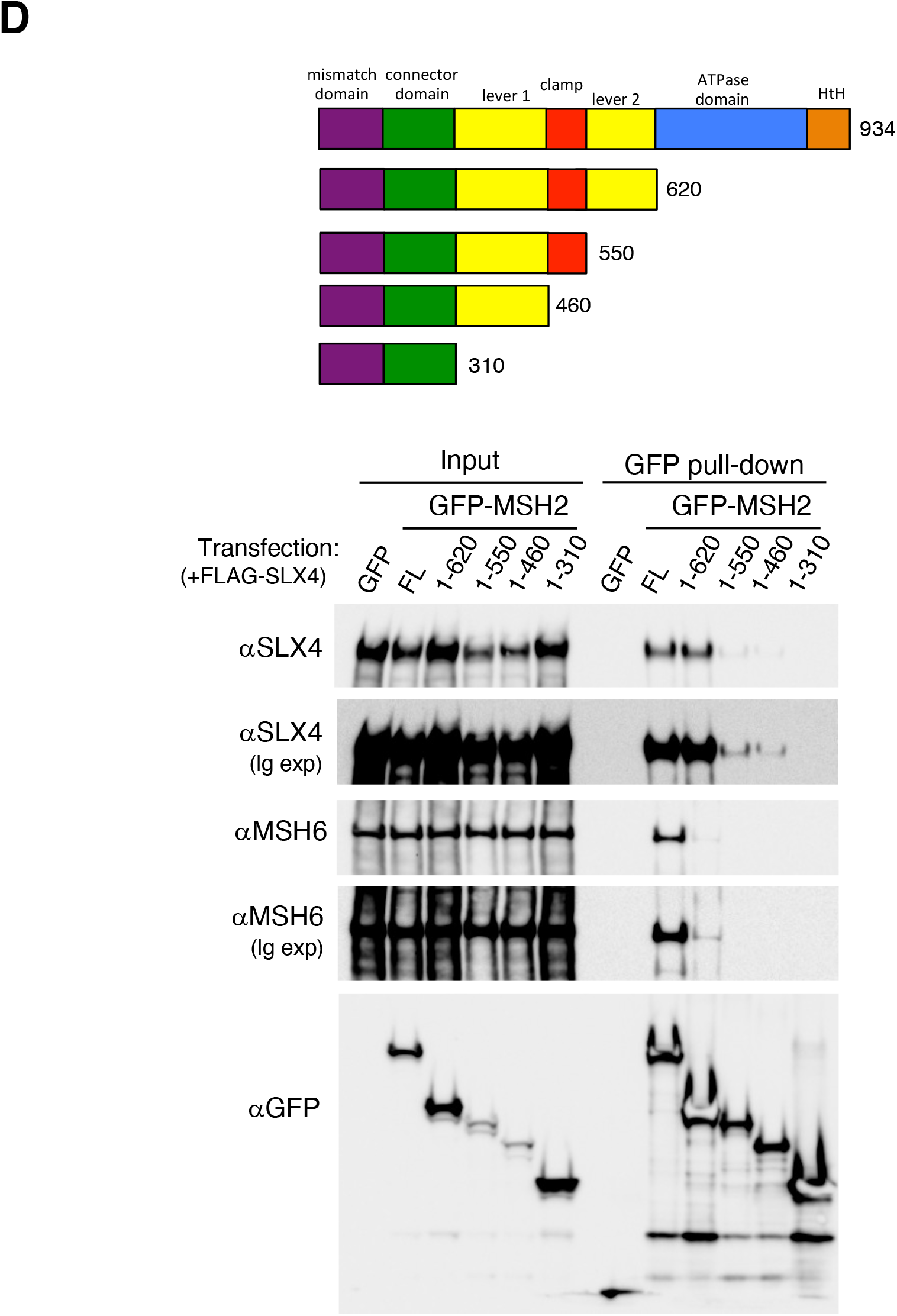
**related to Fig 1**. SLX4 interaction with MSH2. (**A**) Number of peptide spectrum matches (PSM) of MSH2 and MSH6 found in endogenous SLX4 immunoprecipitates using whole cell extracts from control or SLX4-depleted HeLa FITo cells. (**B**) Interaction of MSH2 with the N-terminus of SLX4. The scheme indicates the various FLAG-tagged fragments of SLX4 transfected in HeLa FITo cells and immunoprecipitated before WB analysis. n/a indicates a non-relevant SLX4 mutant used for another research project. (**C**) SLX4 UBZ domains are dispensable for MSH2 interaction. HeLa FITo cells were transiently transfected with YFP-SLX4 WT or a UBZ mutant before GFP pull-down. The asterisk represents an aspecific band recognized by the anti-XPF antibody. (**D**) The lever 1 domain of MSH2 is required for SLX4 interaction. HeLa FITo cells were transfected with FLAG-SLX4 together with the indicated GFP-MSH2 constructs before GFP trap affinity purification and western blotting.

**Figure S2.**
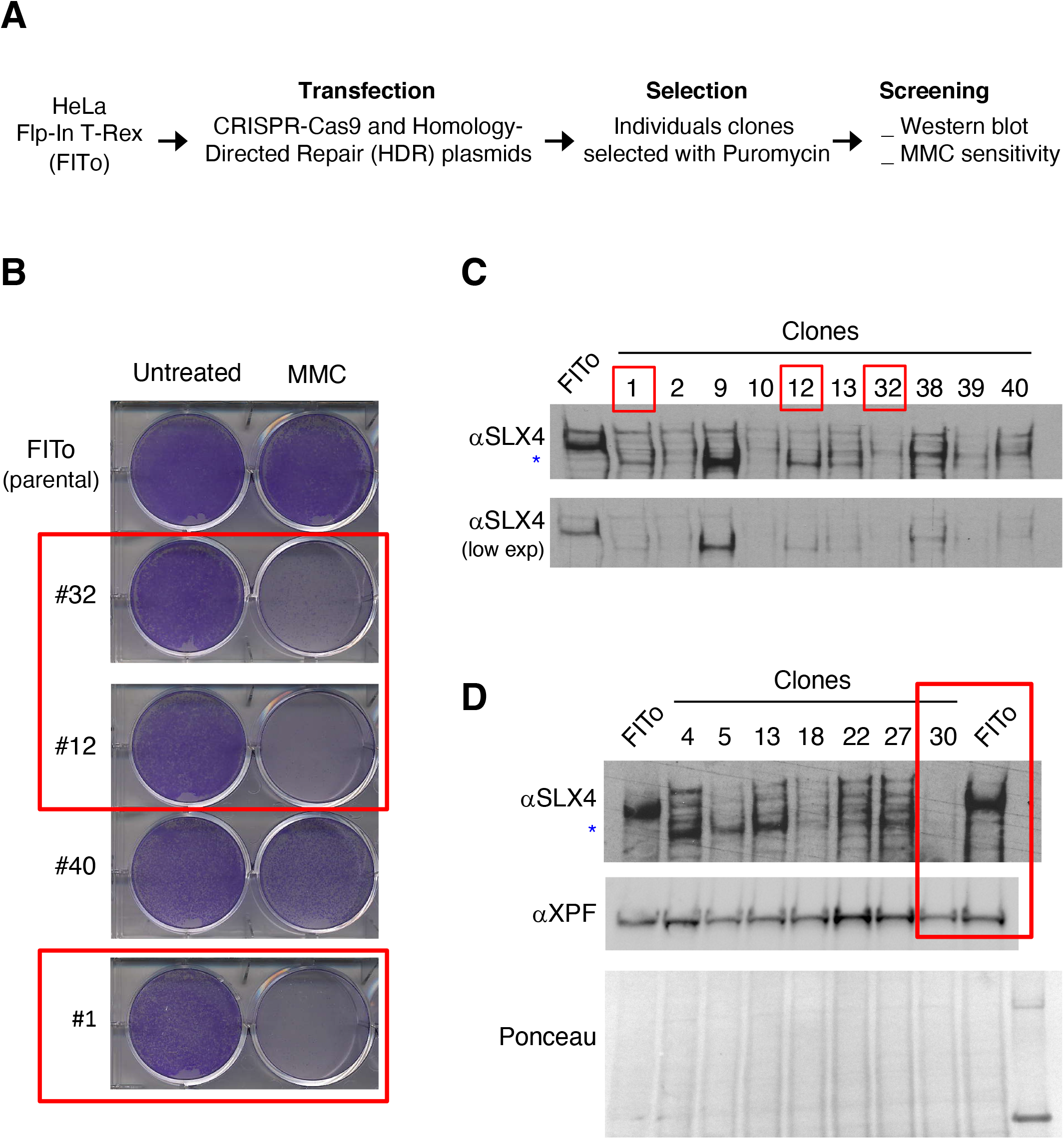
**related to Fig 2**. CRISPR-Cas9 inactivation of SLX4 (**A**) Experimental procedure aiming at knocking out *SLX4* in HeLa cells using CRISPR-Cas9 and homology-directed repair allowing the insertion of a Puromycin-containing plasmid at the *SLX4* locus. (**B**) MMC sensitivity assay of selected Puromycin-resistant clones. Cells were seeded in 6 well plates, treated with MMC (5 ng/ml for 24 h) before drug wash out and addition of fresh medium. Cells were fixed at day 5. The red rectangles indicate clones displaying a severe MMC hypersensitivity. (**C**) and (**D**) Western blots of SLX4 in selected Puromycin-resistant clones using an anti-SLX4 recognizing a C-terminal epitope. The blue asterisk indicates a recurrent band with a lower molecular weight (MW) found in several clones. One clone (KO30) initially displayed an apparent knock-out of SLX4 but the same anti-SLX4 reacting band with a lower MW was found in subsequent analysis.

**Figure S3.**
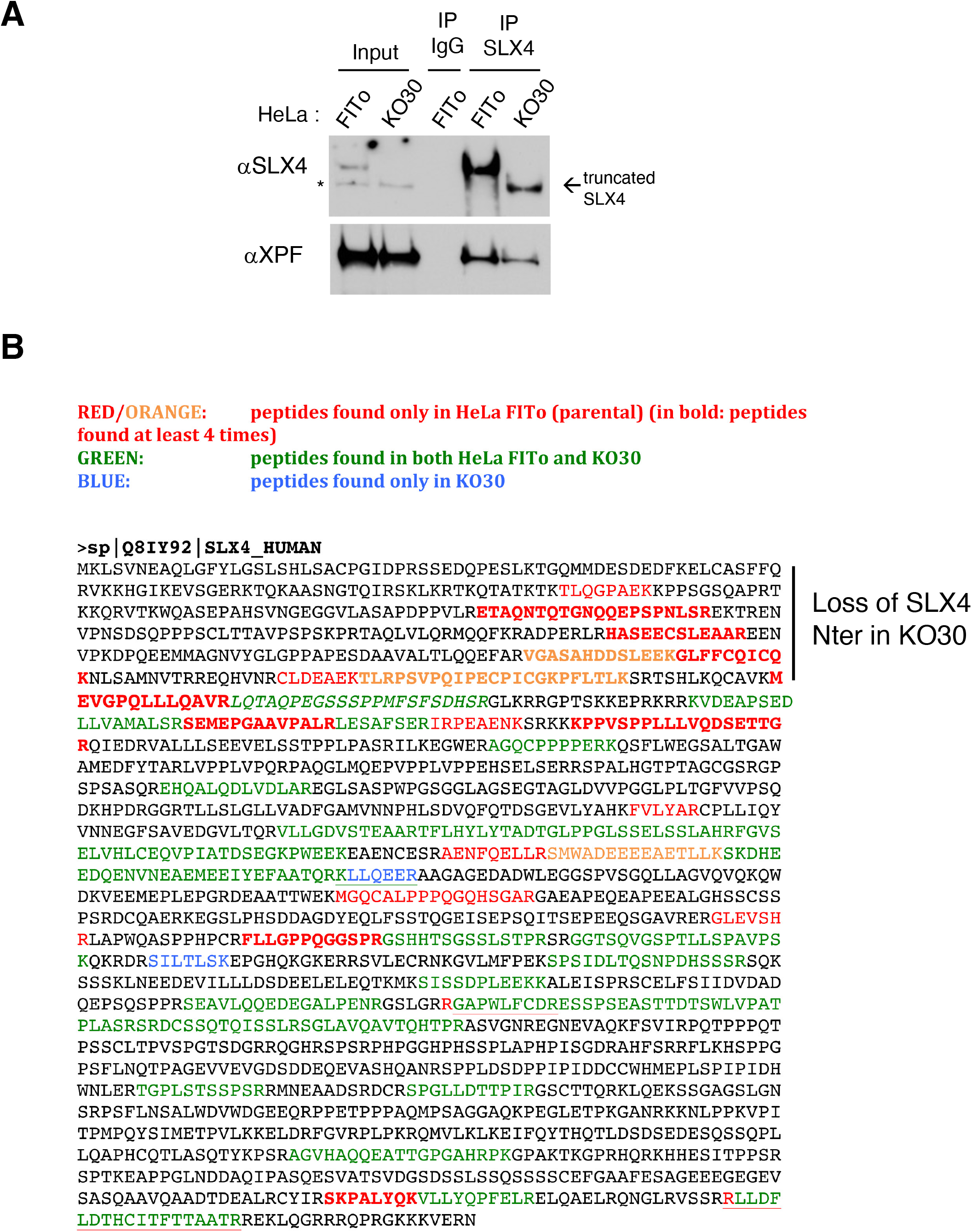
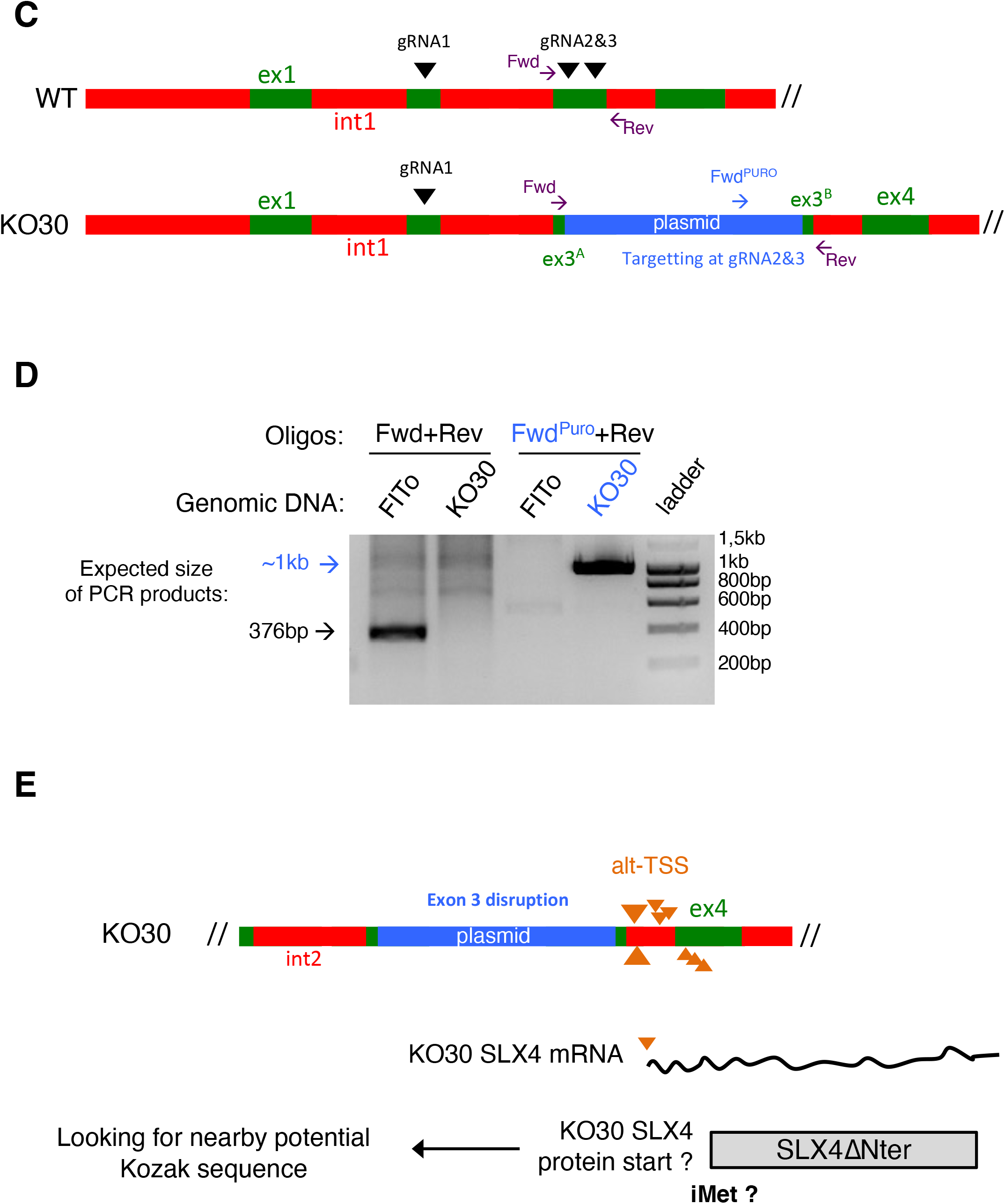
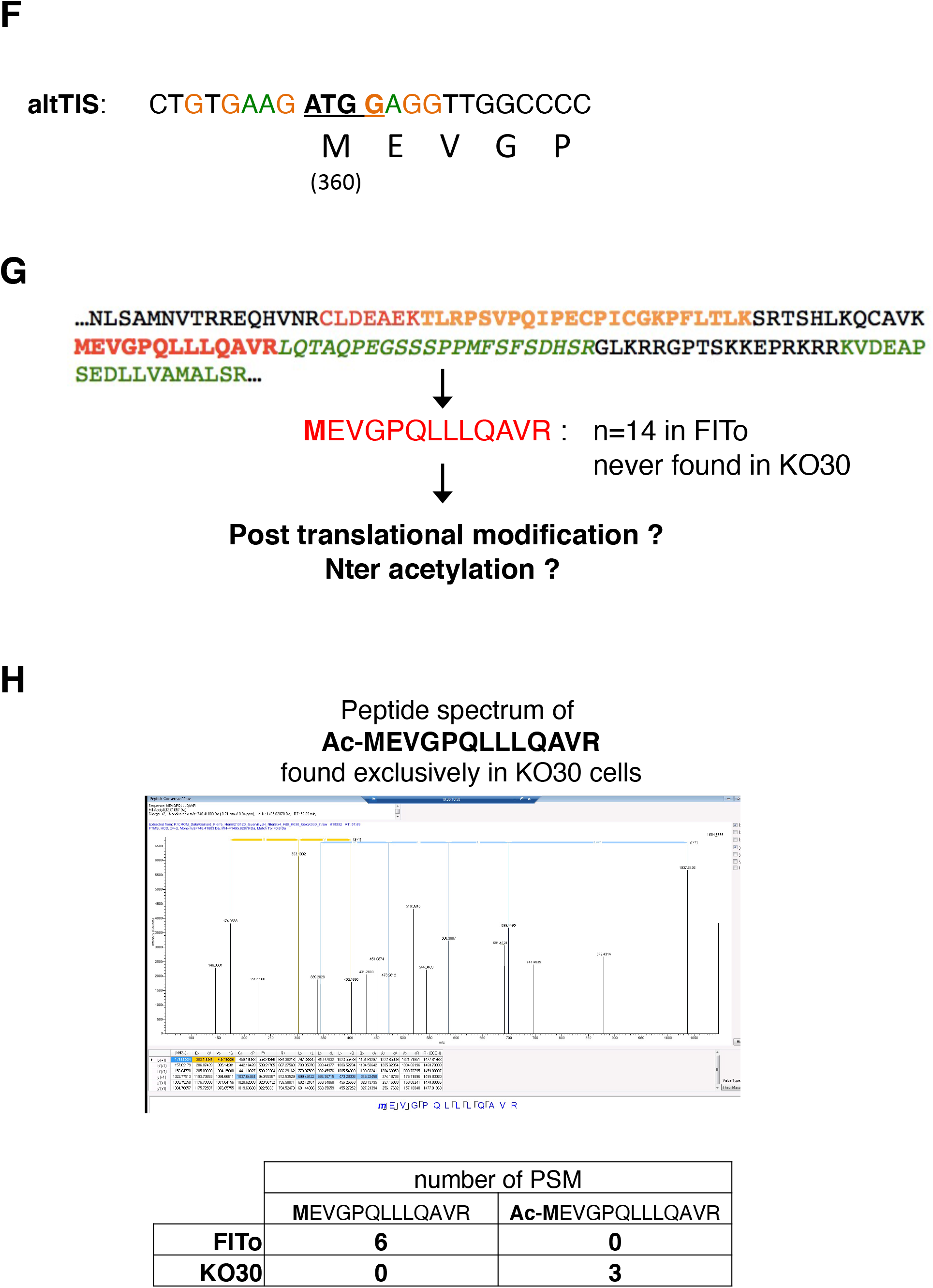
**related to Fig 2**. Characterization of KO30 cells expressing SLX4ΔNter. **(A)** Immunoprecipitation of SLX4 and SLX4ΔNter from FITo and KO30 cells, the asterisk indicates an aspecific band that can be detected by the anti-SLX4 antibody in FITo input and migrates at the same size of SLX4ΔNter but that is not immunoprecipitated. (**B**) Lack of SLX4 N-terminus in KO30 cells. SLX4 immunoprecipitates from HeLa FITo or KO30 cells were analysed by mass spectrometry (MS). As indicated, peptides in red or orange found only in parental FITo cells were overrepresented in SLX4 N-terminus. (**C**) Scheme of the SLX4 locus in FITo or in CRISPR-Cas9 targeted KO30 cells. PCR primers used in (D) are indicated. (**D**) Disruption of Exon 3 integrity and plasmid insertion in KO30 cells. PCR analysis of genomic DNA from FITo and KO30 cells were performed with the indicated primers. (**E**) Model for the generation of SLX4ΔNter in KO30 cells. Alternative Transcription Start sites (Alt-TSS) were found by TSS-seq in several cell lines and reported in the DBTSS (DataBase of Transcriptional Start Sites: https://dbtss.hgc.jp/). Large triangles represent a strong cluster of TSS in the beginning of intron 3 while small triangles represent other TSS in the end of intron 3 and beginning of exon 4. These alt-TSS are compatible with the N-terminal proximal SLX4 peptides found in KO30 cells. (**F**) Candidate alternative Translation Initiation Site (alt-TIS) matching the Kozac sequence consensus. Coloured bases are compatible with a Kozac sequence consensus (1). (**G**) This alt-TIS used in KO30 cells would generate the indicated tryptic peptide that was however found only in FITo cells, possibly because it is post-translationally modified (see also supplementary results). (**H**) Peptide spectrum of the proximal peptide containing an acetylated initiator Methionine found only in KO30 cells after immunoprecipitation of endogenous SLX4 from HeLa FITo or KO30 nuclear extracts and mass spectrometry analysis.

**Figure S4.**
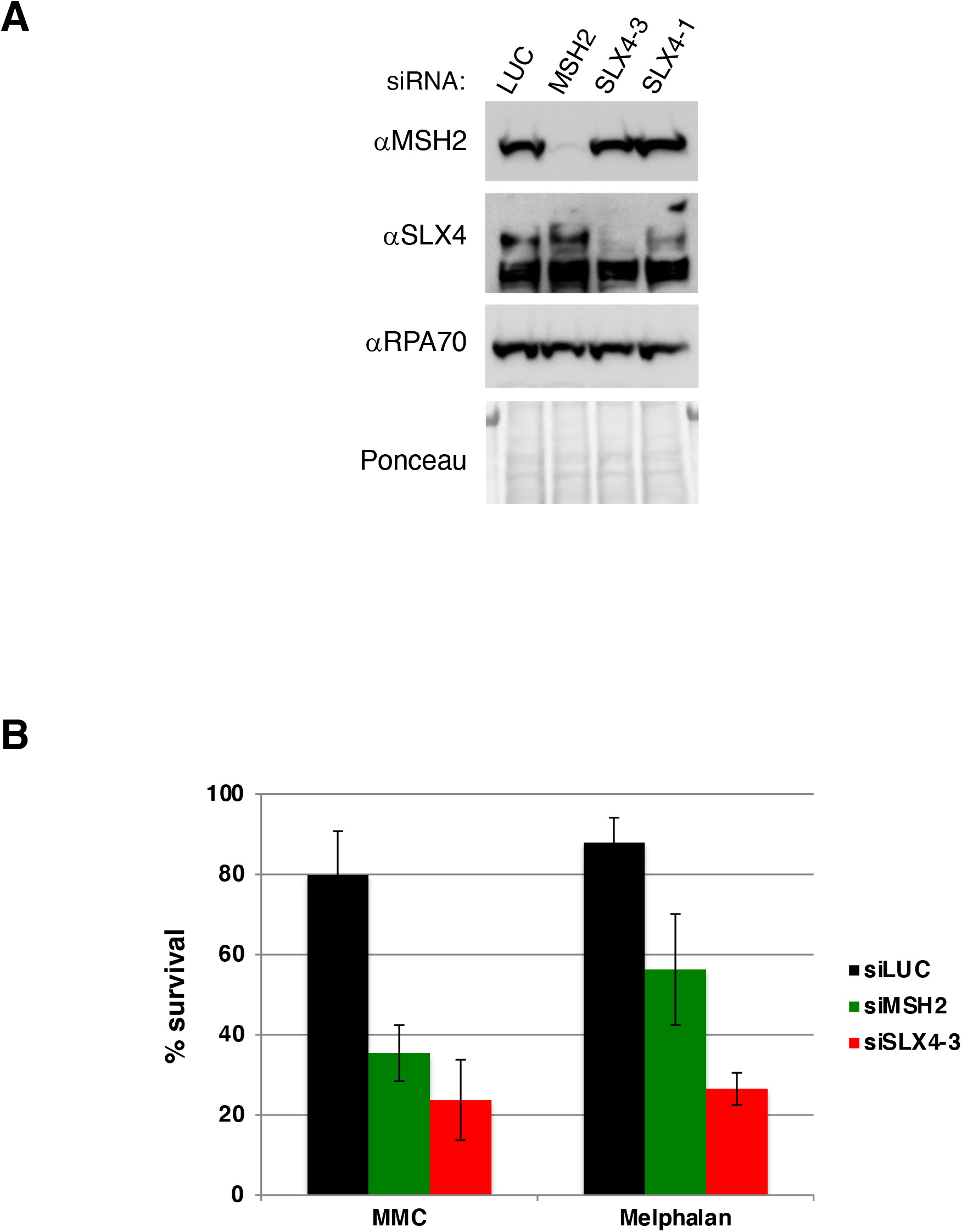
**related to Fig 3 and Fig 4**. (**A**) Western blot showing a representative result of MSH2 or SLX4 depletion in HeLa FITo cells transfected with the indicated siRNAs in clonogenic survival assays. (**B**) Clonogenic survival in response to MMC (5 ng/ml) or Melphalan (500 nM) upon MSH2 or SLX4 depletion (n=3 to 5 experiments, mean ± SD are indicated)

**Figure S5.**
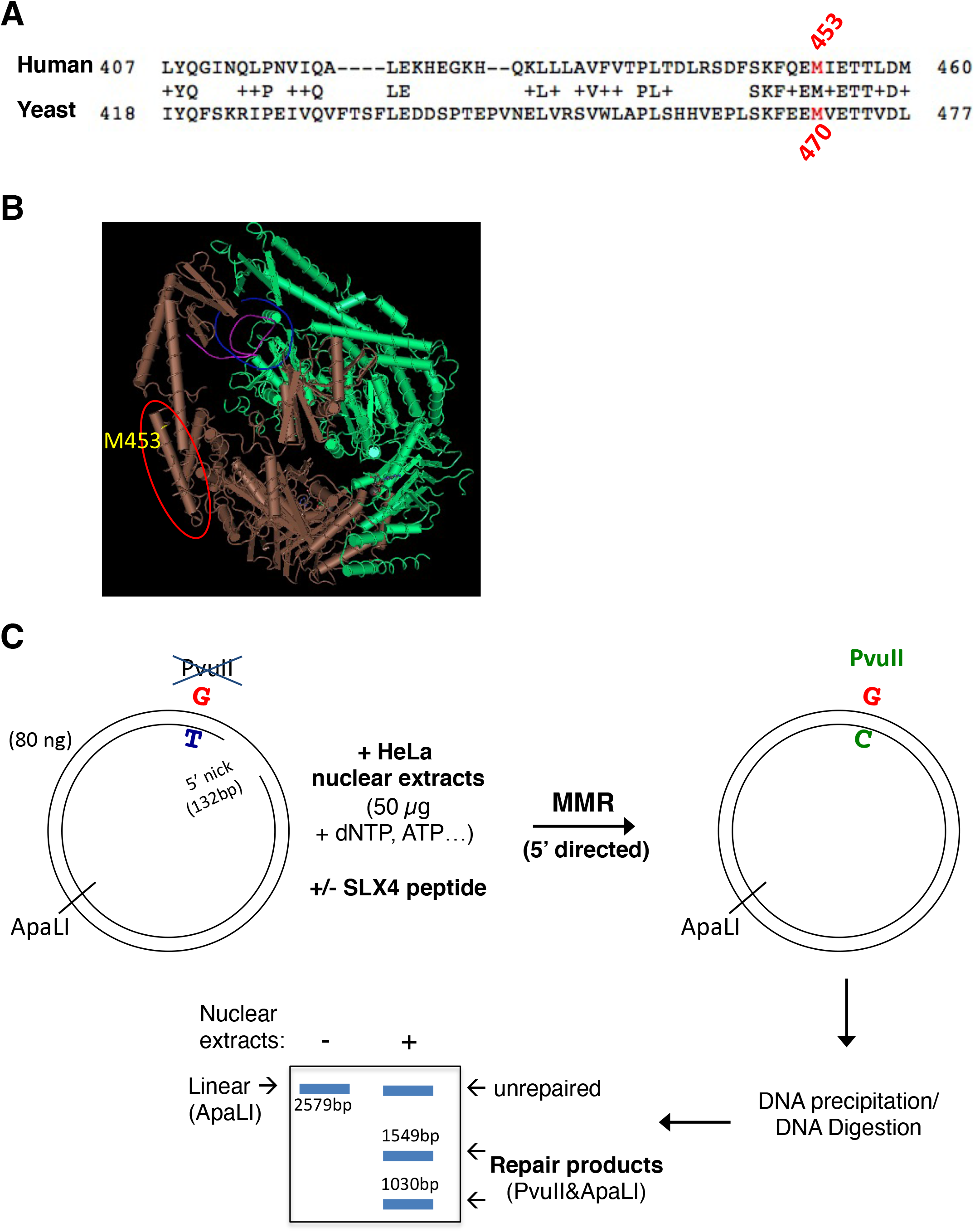
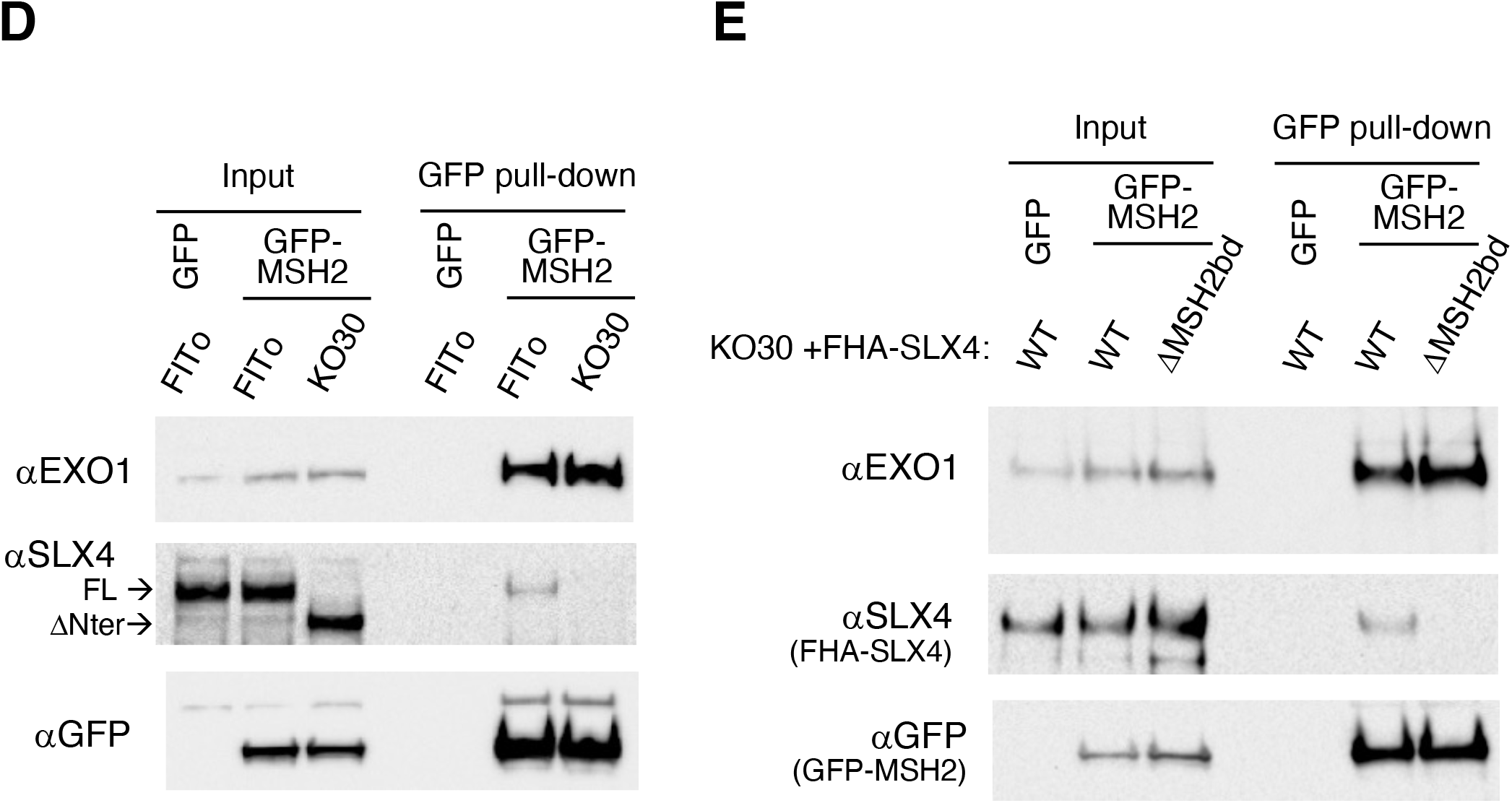
**related to Fig 5**. SLX4 interacts with MSH2 through a SHIP box. (**A**) Alignment of human MSH2 and *S. cerevisiae* Msh2 proteins showing the conservation of M453/M470 and its surrounding residues. (**B**) Structure of the MSH2-MSH6 complex, the position of MSH2 M453 residue is highlighted in yellow. This structure 2O8C represents “ Human MutSα bound to ADP and an O6meG-T mispair” and comes from Warren JJ *et al*. (4). (**C**) Scheme of the *in vitro* MMR assay and example of an expected result, see Material and Methods for details. (**D**) Immunoprecipitation of GFP-MSH2 overexpressed in HeLa FITo or KO30 cells. (**E**) Immunoprecipitation of GFP-MSH2 overexpressed in KO30 cells complemented with FHA-SLX4 WT or FHA-SLX4ΔMSH2bd. Expression of exogenous SLX4 was achieved with 10 ng/ml doxycycline.

